# AGP-Ca^2+^ binding is essential for pollen development and pollen tube growth in Arabidopsis thaliana

**DOI:** 10.64898/2026.01.27.701722

**Authors:** Jessy Silva, Maria João Ferreira, Paul Dupree, Matthew R. Tucker, Maria Manuela Ribeiro Costa, Sílvia Coimbra

**Affiliations:** LAQV/REQUIMTE, Department of Biology, Faculty of Sciences, University of Porto, Porto, 4169-007, Portugal; Department of Biology, University of Minho, Campus of Gualtar, Braga, 4710-057, Portugal; Department of Biochemistry, University of Cambridge, Cambridge, CB2 1QW, UK; Waite Research Institute, School of Agriculture, Food and Wine, The University of Adelaide, Urrbrae, SA, 5064, Australia; Centre of Molecular and Environmental Biology (CBMA), Department of Biology, University of Minho, Campus of Gualtar, Braga, 4710-057, Portugal

## Abstract

Arabinogalactan-proteins (AGPs) are highly glycosylated cell wall proteins essential for plant reproduction, although their mode of action remains unclear. AGPs have been proposed to bind and store calcium (Ca^2+^) in the apoplast via glucuronic acid (GlcA) residues added by glucuronosyltransferases (GLCATs), potentially acting as Ca^2+^ capacitors. Here we report that Ca^2+^ binding by AGPs is required for successful double fertilisation in *Arabidopsis*. Analysis of *glcat14a glcat14b glcat14d* triple mutants revealed reduced seed set due to the abortion of pollen grains, which lacked cytoplasmic content and an intine layer, as confirmed by the absence of cellulose, resembling the phenotype of AGP-deficient mutants. When grown under Ca^2+^-deficient conditions, GLCAT mutants showed exacerbated and conditional reproductive defects that were attenuated under standard Ca^2+^ conditions. Our findings establish a functional requirement for GlcA-mediated Ca^2+^ binding by AGPs and support their role as apoplastic Ca^2+^ stores essential for male fertility and reproductive success in plants.

## Introduction

Double fertilisation is a defining feature of angiosperms that initiates seed formation, involving pollen tube (PT) growth along the female tissues. The male gametophyte or pollen grain (PG) is released from the anthers and adheres to the stigma, where it germinates to form a PT that delivers two male sperm cells to the female gametophyte. During double fertilisation, one male sperm cell fuses with the egg cell, forming the zygote, while the other fertilises the central cell, generating the endosperm, which together will develop into a seed^1^.

In *Arabidopsis thaliana*, PGs develop within the anther locules, surrounded by four somatic single-cell layers: the secretory tapetum, providing materials for pollen wall formation; the middle layer, involved in signalling and nutrient support; the endothecium, required for anther dehiscence; and the outer epidermis, enclosing the anther. Proper coordination and timely degeneration of these layers are essential for pollen wall formation, viability, and dispersal, making them critical for male fertility^2^.

Pollen formation begins with microsporogenesis, during which diploid pollen mother cells undergo meiosis to produce tetrads of haploid microspores within a callose wall, later released by β-1,3-glucanase secreted from the tapetum. During microgametogenesis, the released uninucleate microspores enlarge and divide asymmetrically (pollen mitosis I) to form a bicellular pollen grain consisting of a vegetative cell and a generative cell. The generative cell then divides (pollen mitosis II) to produce two sperm cells, which, together with the vegetative nucleus, constitute the male germ unit, a functional unit essential for fertilisation. The pollen wall is a specialised extracellular multilayer that protects PGs, facilitates pollination, and mediates pollen-pistil interactions. It consists of three layers: the inner intine, rich in pectins, cellulose, and hemicelluloses secreted by the microspores; the exine, composed of sporopollenin derived from tapetal secretions; and the pollen coat, formed from tapetal remnants after programmed cell death. The intine includes a granular exintine and microfibrillar endintine sublayers, while the exine comprises an inner bilayer nexine and an outer reticulated sexine sublayers containing the tectum and bacula^3–5^.

Arabinogalactan-proteins (AGPs) are involved in multiple stages of plant reproduction, including gametophyte development, male-female interactions, and seed development^3,6,7^. AGPs are among the most complex and diverse families of glycoproteins in plants, extensively decorated with arabinogalactan (AG) polysaccharides — composed mainly of arabinose and galactose, with minor sugars such as glucuronic acid (GlcA), fucose, and rhamnose — which can constitute up to 90% of the molecule. Some AGPs are anchored to the plasma membrane via a glycosylphosphatidylinositol (GPI) lipid anchor^8^. Despite their importance, elucidating AGP function through genetic approaches has been challenging due to genetic redundancy^9^.

A longstanding hypothesis proposes that AGPs bind and store calcium ions (Ca^2+^) through GlcA residues in the apoplast^10^. Ca^2+^ signalling is essential for plant reproduction, with distinct and regulated spatiotemporal Ca^2+^ signatures occurring as the PT grows along the stigma and style, interacts with the synergid cell and during gamete activation and fusion^1,11^. Experimental evidence further supports a direct role for AGPs in Ca^2+^ regulation as β-Yariv binding to AG polysaccharides increases intracellular Ca^2+^ levels^12^ and inhibits PT growth^13^, implicating AGPs and their glycans in reproductive Ca^2+^ dynamics^14^. Consequently, some AGPs expressed in reproductive tissues may function as surface Ca^2+^ capacitors.

AGP glycosylation is mediated by a network of 24 identified glycosyltransferases (GTs) in *Arabidopsis*, including galactosyltransferases (GALTs), arabinosyltransferases, fucosyltransferases, rhamnosyltransferases, xylosyltransferases, glucuronosyltransferases (GLCATs), and GlcA methyltransferases^8,15,16^. GLCATs, responsible for adding GlcA to AGPs, belong to the CAZy GT14 family, which comprises 11 members in *Arabidopsis*^8,17^. Five of these (GLCAT14A, GLCAT14B, GLCAT14C, GLCAT14E, and GLCAT14F) have been characterised as β-1,6-GLCATs^16,18,19^, while GLCAT14D has been shown to be essential for AG glucuronidation^20^. In a previous study, the *glcat14a glcat14b glcat14d-1* triple mutant contained 66% less methylated GlcA on leaf AGs and bound approximately 80% less Ca^2+^ than wild-type (WT) plants, validating the hypothesis of AGP-Ca^2+^ interaction via GlcA^20^.

Here, we demonstrate that Ca^2+^ binding by glucuronidated AGPs is essential for male reproductive success in *Arabidopsis*. Analysis of *glcat14a glcat14b glcat14d* triple mutants revealed severe defects in male gametophyte development, including collapsed PGs devoid of cytoplasmic content and intine layer. These defects led to reduced pollen viability, lower germination rates, and shortened PTs, ultimately causing a marked decrease in seed set. The phenotypes were intensified under low Ca^2+^ conditions and were partially alleviated under Ca^2+^-sufficient conditions. Notably, the mutant defects resemble those observed in AGP-deficient plants, supporting a model in which GlcA-mediated Ca^2+^ binding by AGPs serves as an apoplastic Ca^2+^ reservoir essential for pollen development, PT growth, and successful double fertilisation.

## Results

### *GLCAT14A/B/D* triple mutants show reduced growth and fertility

To elucidate the contribution of GlcA to plant sexual reproduction, we characterised two previously reported triple mutants, *glcat14a-2 glcat14b-1 glcat14d-1* and *glcat14a-2 glcat14b-1 glcat14d-2*^20^, hereafter referred to as *glcat14a/b/d-1* and *glcat14a/b/d-2*, respectively. Both lines carry T-DNA insertions disrupting the coding regions of *GLCAT14A*, *GLCAT14B*, and *GLCAT14D*, and reverse transcription quantitative real-time PCR (RT-qPCR) revealed null expression in *glcat14a/b/d-1* and *glcat14a/b/d-2* mutants compared to WT plants, confirming that both lines were triple knockout mutants (Extended Data Fig. 1a,b). Mutant plants exhibited delayed growth and reduced stature relative to WT (Extended Data Fig. 1c), with stem length reduced by 18–27% (Extended Data Fig. 1d) and fewer developed siliques (Extended Data Fig. 1e). A summary of growth and reproductive phenotypes of *glcat14a/b/d* mutants is provided in Supplementary Table 1. Despite these growth and fertility defects, the overall arrangement and morphology of floral organs from stages 12–15^21,22^ appeared indistinguishable from WT, with normal sepals, petals, pistils, and anthers (Extended Data Fig. 2a). Quantitative analysis of stamen and pistil lengths across the same flower stages confirmed that organ growth was comparable between WT and mutants at most stages. At flower stage 13, *glcat14a/b/d-2* showed slightly reduced stamen and pistil lengths relative to WT, and at stage 14, stamen length remained decreased (Extended Data Fig. 2b,c). Notably, many flowers in both triple mutants failed successful pollination (Extended Data Fig. 2a, right column), despite normal organ size and positioning, suggesting that loss of *GLCAT14A/B/D* expression can compromise reproductive success through effects on pollen release, viability, or stigma receptivity.

### Seed set defects in *glcat14a/b/d* mutants

To investigate reproductive defects, we analysed siliques at flower stage 17^21,22^, and both triple mutants exhibited seedless regions within their siliques (Fig. 1a). Silique length was reduced by 16–19% in *glcat14a/b/d* mutants compared with WT (Fig. 1), and seed set was highly variable, showing a significant 14–18% decrease relative to WT (Fig. 1c). To determine whether the reproductive defects originated from male or female gametophytes, we performed self- and reciprocal crosses between WT and the triple mutants. Low pollen release in the mutants made manual crosses more laborious, often requiring pollen from multiple flowers. Cleared siliques at 10 days after pollination (DAP) revealed seedless regions in all reciprocal combinations (Fig 1d). Silique length and seed set were reduced by 10–25 % and 16–31 %, respectively, in all mutant crosses compared to WT self-cross (Fig 1e,f), though not statistically significant. No difference was observed between self- and reciprocal crosses of *glcat14a/b/d* mutants, suggesting that defects are not restricted to a single gametophyte and may involve sporophytic contributions.

**Fig 1.**
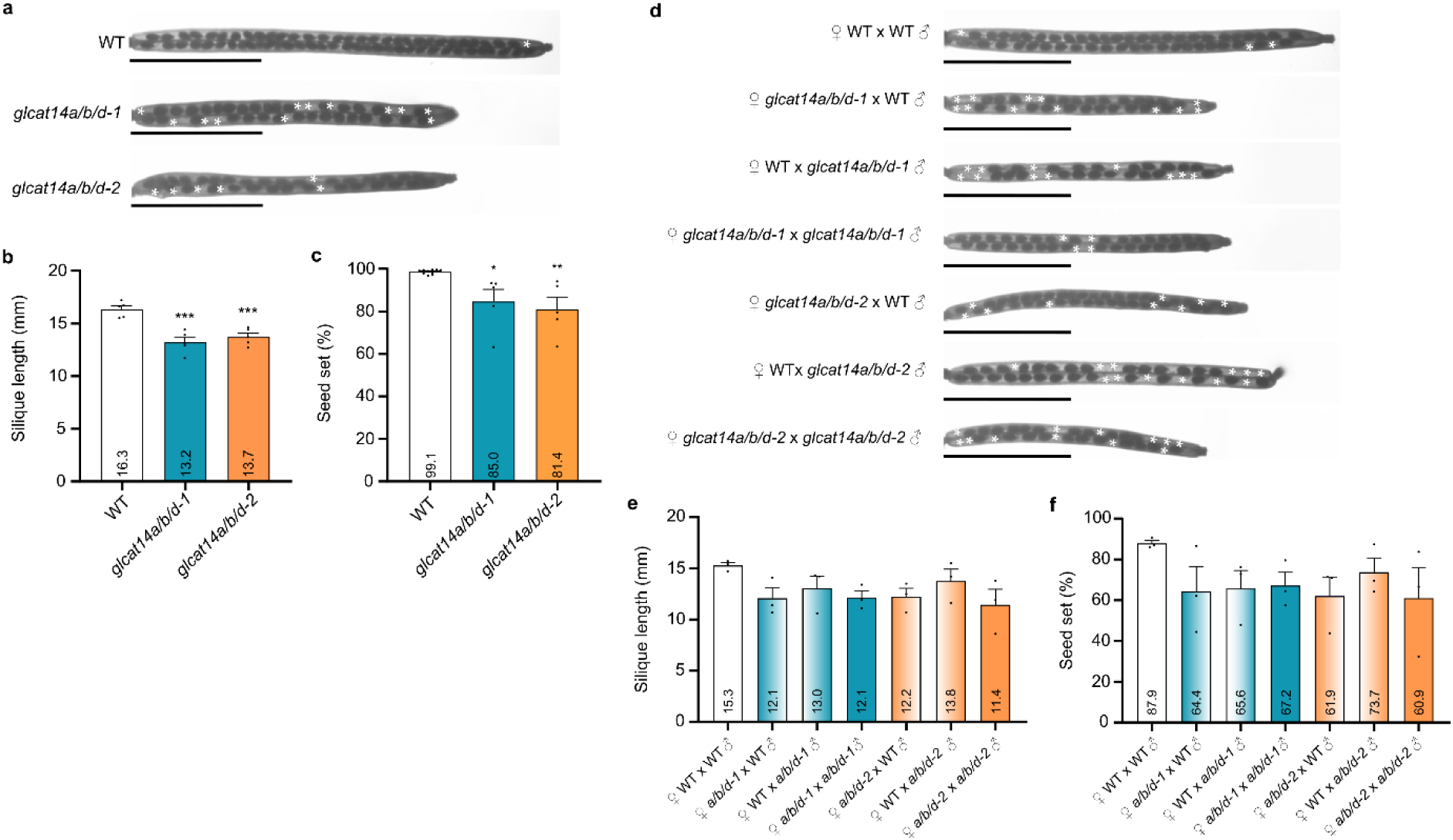
Reduced fertility in *glcat14a/b/d* mutants. **a**, Cleared siliques from seven-week-old wild-type (WT), *glcat14a/b/d-1*, and *glcat14a/b/d-2* plants. **b**,**c**, Silique length (**b**) and seed set percentage (**c**) of seven-week-old WT and *glcat14a/b/d* mutants. Data are presented as mean + SEM (**b**: *n* = 5 plants, 5 siliques per plant; **c**: *n* = 5 plants per *glcat14abd* allele and *n* = 10 plants for WT, 10 siliques per plant). **d**, Cleared siliques 10 days after pollination (DAP) from hand-pollinated self and reciprocal crosses between WT and *glcat14a/b/d* mutants. **a**,**d**, Areas without seeds are marked with white asterisks. Scale bars, 5 mm. **e**,**f**, Silique length (**e**) and seed set percentage (**f**) of siliques 10 DAP from hand-pollinated crosses between WT and *glcat14a/b/d* mutants. Data are presented as mean + SEM (*n* = 3 plants/crosses, 5 siliques per plant). Biological replicates are shown as black dots. Mean values are shown within bars. Statistical significance was determined by one-way ANOVA followed by Dunnett’s test (**b**,**c**) or by Tukey’s test (**e**,**f**). Asterisks represent statistically significant differences from WT (*, *p*<0.0332; **, *p*<0.0021; ***, *p*<0.0002). No significant differences were observed in (**e**) and (**f**).

### Pollen grain collapse and reduced pollen tube growth in the mutants

To assess whether fertility loss may be due to defects in male gametopyhte development, we next assessed pollen viability using Alexander staining. At flower stage 12, *glcat14a/b/d-1* and *glcat14a/b/d-2* mutant anthers contained mostly red-magenta PGs, indicative of viability, alongside a subset of green-stained non-viable PGs (Fig. 2a). After anthesis at flower stage 13, PGs remained heterogeneous, with the majority resembling WT and showing the characteristic rounded red-magenta staining of viable PGs, wheareas a substantial fraction appeared shrunken and greenish, consistent with aborted PGs (Fig. 2a). Quantitative analysis confirmed that approximately 32% of PGs were aborted in both *glcat14a/b/d* mutants compared to nearly 100% viability in WT (Fig. 2b). Staining of anthers with decolourised aniline blue solution (DABS) revealed autofluorescent pollen walls and highlighted collapsed PGs in the mutants (Fig. 2c). DAPI staining confirmed the presence of sperm and vegetative nuclei in viable PGs of all genotypes, whereas shrunken PGs in mutants lacked nuclei (Fig. 2d).

**Fig 2.**
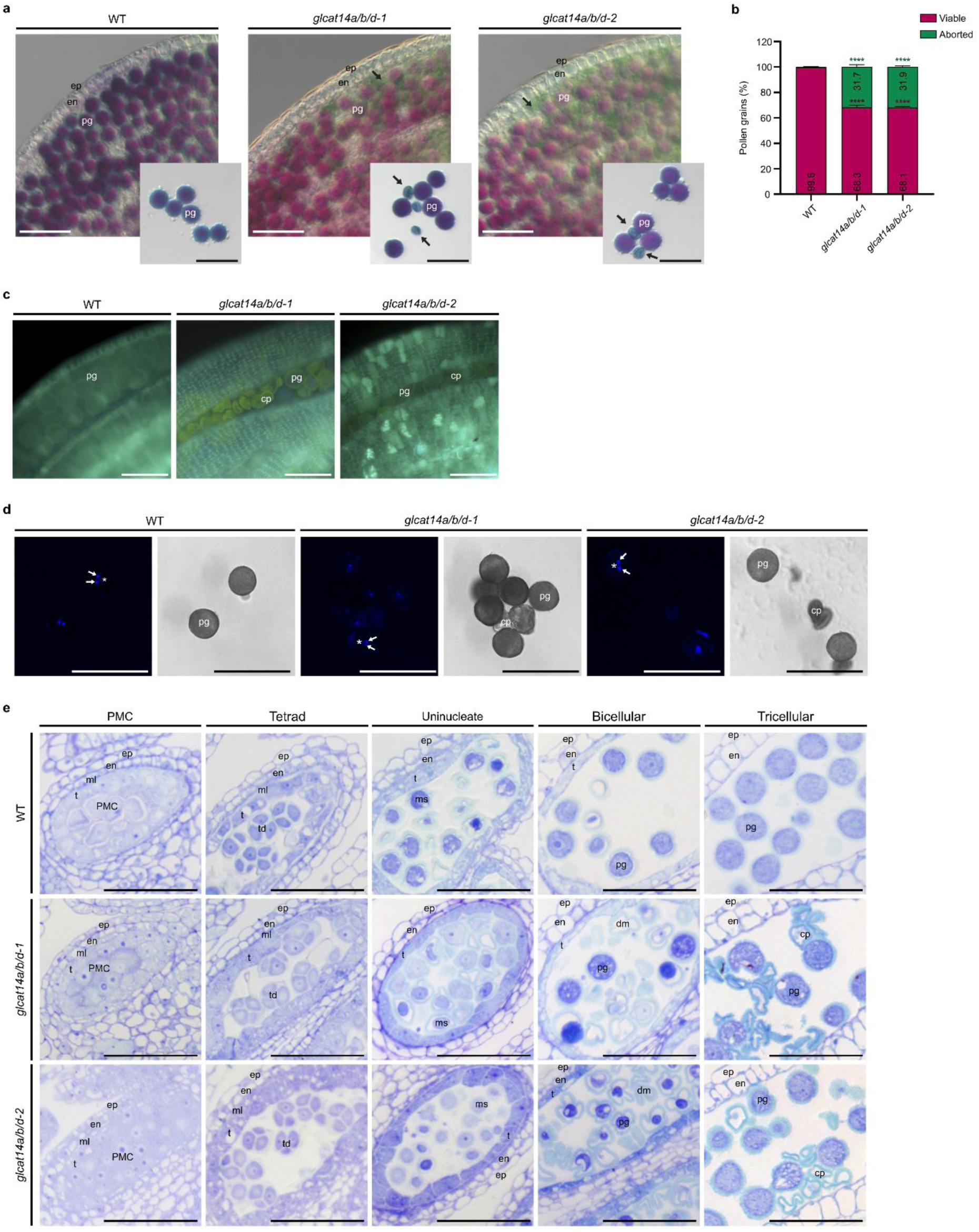
Pollen grain collapse and impaired development in *glcat14a/b/d* mutants. **a**, Alexander staining of anthers at flower stage 12 and pollen grains (PGs) of anthers at flower stage 13 of wild-type (WT), *glcat14a/b/d-1*, and *glcat14a/b/d-2* plants. Viable PGs are stained in red-magenta while aborted PGs are stained in blue-green (black arrows). **b**, Viable and aborted PGs percentage of WT and *glcat14a/b/d* mutants. Data are presented as mean + SEM (*n* = 3 plants, 500 PGs per plant). Mean values are shown within bars. Statistical significance was determined by two-way ANOVA followed by Dunnett’s test. Asterisks represent statistically significant differences from WT (****, *p*<0.0001). **c**, Aniline blue staining of anthers at flower stage 13 of WT and *glcat14a/b/d* plants. **d**, DAPI staining of PGs of anthers at flower stage 13 of WT and *glcat14a/b/d* mutants. Sperm cell (white arrows) and vegetative (white asterisks) nuclei are shown in the DAPI channel, alongside the corresponding brightfield channel indicating PG boundaries. **e**, Anther sections at flower stages 9–12, showing pollen mother cell, tetrad, uninucleate microspore, bicellular, and tricellular PG stages of WT and *glcat14a/b/d* plants. Abbreviations: cp, collapsed pollen grain; dm, degenerated microspore; en, endothecium; ep, epidermis; ml, middle layer; ms, microspore; pg, pollen grain; PMC, pollen mother cell; t, tapetum; td, tetrad. Images were acquired using a Zeiss Axio Imager.A2 microscope with DIC optics (**a**,**e**), Leica DMLB epifluorescence microscope (**c**), and Leica TCS SP5 II confocal system (**d**). Scale bars, 50 μm.

To determine the timing of pollen abortion, anthers at flower stages 9 to 12^21,22^ were sectioned and stained with toluidine blue. At the PMC and tetrad stages, tapetum and locules of mutants were indistinguishable from WT, with no observable abnormalities (Fig. 2e). At the uninucleate stage, microspores in mutants became vacuolated, similar to WT. However, by the bicellular stage, nearly half of mutant PGs collapsed, in contrast to normal development in WT. Tapetum degeneration proceeded normally in mutants during the late stages of pollen development. By the tricellular stage, mutant locules contained collapsed PGs devoid of cytoplasm and surrounded by exine remnants, which were absent in WT (Fig. 2e). These results indicate that pollen abortion in *glcat14a/b/d* mutants occurs between the uninucleate and bicellular stages.

### Pollen wall defects and reduced pollen performance in *glcat14a/b/d* mutants

To investigate the mechanism of PG abortion in *glcat14a/b/d* mutants, we analysed the pollen wall at flower stage 12. Autofluorescence imaging of anther sections, which highlights the sporopollenin-rich exine, revealed that while the exine layer was present in both viable and collapsed PGs, it was distorted and shrunken in the collapsed PGs (Fig. 3a). To examine the intine layer, sections were stained with calcofluor white, which binds β-glucans including cellulose and callose, and with DABS, which specifically labels callose. Collapsed PGs showed no staining with either marker, indicating a complete loss of β-glucans and callose in the intine, whereas viable PGs displayed a well-defined circular signal corresponding to an intact intine. GFP–CBM3 labelling confirmed the absence of cellulose in collapsed PGs of *glcat14a/b/d* mutants. In viable PGs of these mutants, the GFP–CBM3 signal was irregular in shape, whereas in WT PGs it formed a well-defined circular inner layer beneath the exine, suggesting subtle defects in intine organisation even in PGs that remains cytologically intact (Fig. 3a).

**Fig 3.**
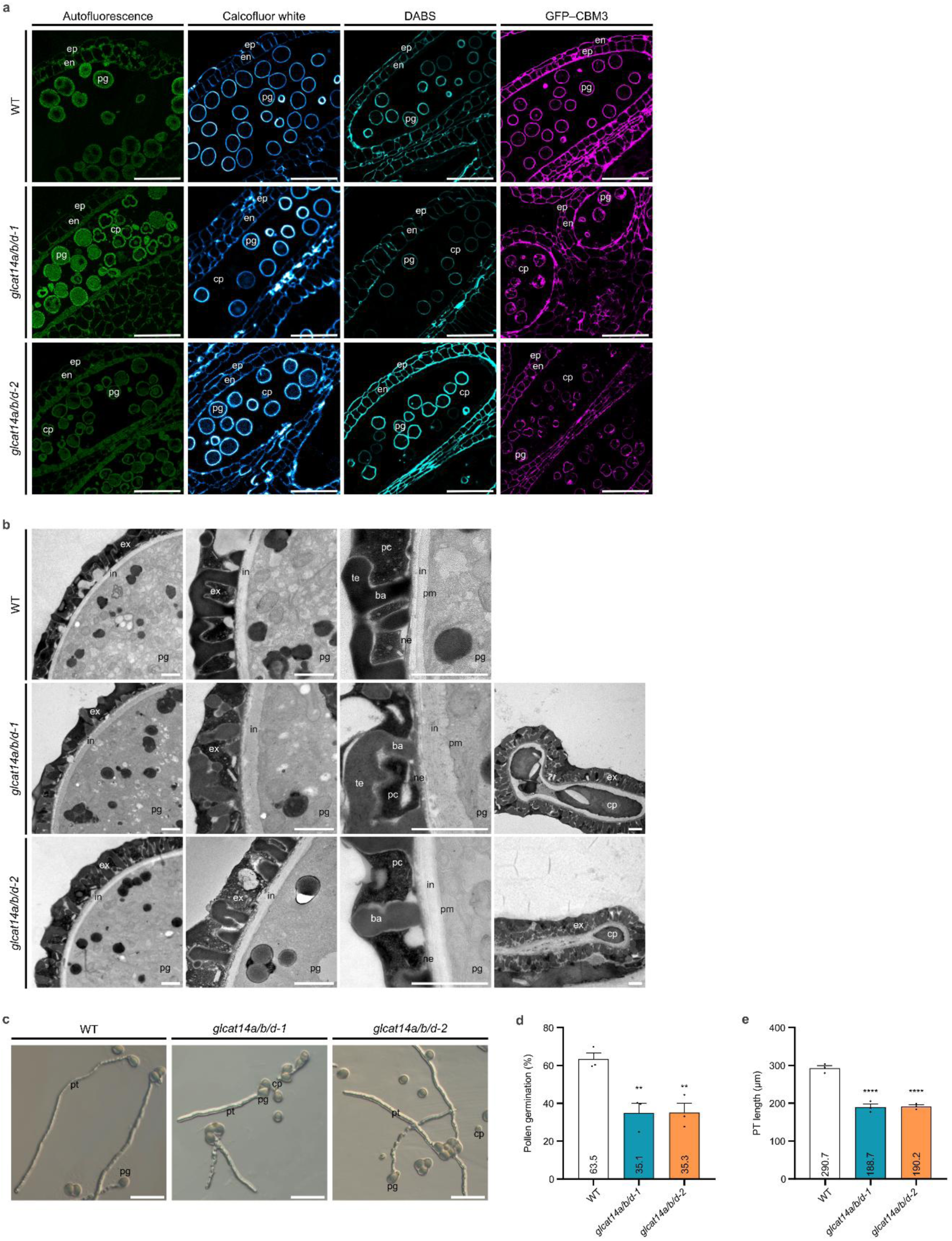
Pollen-wall defects and impaired germination in *glcat14a/b/d* mutants. **a**, Autofluorescence, calcofluor white, decolourised aniline blue solution (DABS), and GFP–CBM3 staining of anther sections at flower stage 12 of wild-type (WT), *glcat14a/b/d-1*, and *glcat14a/b/d-2* plants. **b**, Transmission electron microscopy of pollen grains (PGs) of WT and *glcat14a/b/d* mutans. **c**, *In vitro* pollen germination of PGs of WT and *glcat14a/b/d* plants. Abbreviations: ba, bacula; cp, collapsed pollen grain; en, endothecium; ep, epidermis; ex, exine; in, intine; ne, nexine; pc, pollen coat; pg, pollen grain; pm, plasma membrane; pt, pollen tube; te, tectum. Images were acquired using a Leica TCS SP8 confocal system (**a**), JEOL JEM 1400 transmission electron microscope (**b**), and Zeiss Axio Imager.A2 microscope with DIC optics (**c**). Scale bars, 1 μm (**b**), 50 μm (**a**), and 100 μm (**c**). **d**,**e**, Pollen germination percentage (**d**) and pollen tube (PT) length (**e**) of WT and *glcat14a/b/d* mutants. Data are presented as mean + SEM [*n* = 3 plants, 150 PGs per plant for (**d**) and 50 PTs per plant for (**e**)]. Biological replicates are shown as black dots. Mean values are shown within bars. Statistical significance was determined by one-way ANOVA followed by Dunnett’s test. Asterisks represent statistically significant differences from WT (**, *p*<0.0021; ****, *p*<0.0001).

To complement the staining analyses, we examined pollen ultrastructure by transmission electron microscopy (TEM) of anthers at flower stage 12. Tapetal cells had degenerated and released their contents into the exine cavities to form the pollen coat in both WT and *glcat14a/b/d* mutant anthers (Fig. 3b). In both genotypes, PGs exhibited a well-organised exine with clearly defined bacula, tectum, and nexine. In WT PGs, a continuous intine layer was consistently observed beneath the exine. By comparison, viable PGs in *glcat14a/b/d* mutants displayed an irregular and slightly thickened intine. Collapsed PGs additionally displayed degenerated cytoplasm and a complete absence of the intine layer (Fig. 3b). These observations indicate that loss of GLCAT14A/B/D activity severely compromises pollen wall formation and integrity, with collapsed PGs displaying loss of cytoplasmic integrity and a complete absence of the intine layer, reflecting a defect in the final stages of pollen development.

To determine whether these structural defects impact pollen performance, we next examined pollen germination and PT growth in vitro. Collapsed PGs failed to germinate, confirming that only morphologically viable PGs could produce PTs. Viable PGs from *glcat14a/b/d* mutants germinated at a significantly lower rates than WT, corresponding to a 45% reduction (Fig. 3c,d). PTs of *glcat14a/b/d* mutants were also shorter than WT, representing a 35% reduction in PT length (Fig. 3c,e). Considering that 32% of total PGs are collapsed, only 24% of total PGs in the mutants were able to germinate, and their PTs were markedly shorter. These results indicate that, in addition to reduced pollen viability, surviving viable pollen exhibit impaired germination and PT elongation, suggesting that defective pollen-wall formation compromises pollen performance.

### Altered AGP and pectin deposition in anthers of *glcat14a/b/d* mutants

To investigate AGP distribution and abundance in *glcat14a/b/d* mutants, we performed fluorescent immunolocalisation on anther sections at flower stage 12 using antibodies that recognize AGP epitopes. JIM13 labelled the cell walls of endothecial cells in both WT and mutants (Extended Data Fig. 3). JIM8 strongly labelled the endothecium in WT, while labelling in *glcat14a/b/d* mutants was slightly reduced and more punctate. Sperm cell walls were labelled with both JIM13 and JIM8 in all genotypes. LM2 produced dot-like signals in anther cells of both WT and mutants without a defined pattern and with a slight reduction in signal intensity in the mutants. LM14 labelled the epidermal and endothecial cell walls in WT, with reduced labelling in the mutants, and also marked the intine layer of PGs in all genotypes. Notably, collapsed PGs in *glcat14a/b/d* mutants lacked detectable labelling with any of the AGP antibodies tested (Extended Data Fig. 3). To assess whether intine defects in glcat14a/b/d mutants were accompanied by altered pectin deposition, anther sections were labelled with JIM5 and JIM7. JIM7 produced strong signals in the cell walls of the epidermis and endothecium, as well as in the intine layer of WT pollen, with a similar labelling pattern observed in the mutants. JIM5 revealed weaker and more punctate labelling in the epidermis, endothecium, and intine layer in both WT and mutant anthers. Collapsed PGs in *glcat14a/b/d* mutants were predominantly unlabelled by JIM5 and JIM7, with only occasional weak signals (Extended Data Fig. 3), indicating that pectins are largely absent and supporting that intine formation is severely compromised. Overall, immunolabelling analyses did not reveal a major reduction in AGP epitope labelling in *glcat14a/b/d* anther locules, suggesting that reproductive defects are unlikely due to loss of AGP epitopes, but may instead arise from altered AGP–Ca^2+^ interactions.

### Enhanced growth and reproductive defects under low Ca²⁺ conditions

To investigate whether the reproductive defects in *glcat14a/b/d* mutants result from impaired AGP–Ca^2+^ interactions due to reduced GlcA content and lower Ca^2+^ availability, we assessed fertility under defined Ca^2+^ conditions. Previous studies showed that *glcat14a/b/d* mutants are hypersensitive to low external Ca^2+^, and their developmental phenotypes can be partially rescued by providing normal Ca^2+^ levels^20^. To assess how reduced Ca^2+^ availability affects reproductive development, *glcat14a/b/d-1* and *glcat14a/b/d-2* plants were grown hydroponically in nutrient solutions containing either low (0.2 mM; low Ca^2+^ nutrient solution - LCNS) or basal (2 mM; basal nutrient solution - BNS) Ca^2+^ concentrations. Mutant plants remained smaller than WT at three and four weeks old in both BNS and LCNS (Extended Data Fig. 4a). Flowering time, measured as days to anthesis of the first flower, was delayed in *glcat14a/b/d-1* compared to WT under BNS, and in both *glcat14a/b/d-1* and *glcat14a/b/d-2* compared to WT under LCNS. A slight delay was also observed in WT under LCNS relative to BNS, and in *glcat14a/b/d-1* under LCNS compared with BNS (Extended Data Fig. 4b). Similar to earlier observations on Ca^2+^ sensitivity^20^, at five weeks old, mutant plants grown in BNS remained significantly shorter than WT, and under LCNS, mutants were even smaller, showing a stronger reduction compared not only to WT but also to their BNS-grown counterparts (Extended Data Fig. 4c,d). Plants grown in LCNS developed fewer and shorter siliques than those in BNS. Under BNS, *glcat14a/b/d* mutants produced fewer siliques than WT, a defect further exacerbated under LCNS, where both mutants formed markedly fewer and smaller siliques compared with WT (Extended Data Fig. 4e). Supplementary Table 2 provides a summary of the growth and reproductive defects of *glcat14a/b/d* plants.

Siliques of *glcat14a/b/d* mutants grown with BNS showed partial areas lacking mature seeds, a phenotype that was more pronounced under LCNS (Fig. 4a). Silique length and seed set were significantly reduced in mutants under LCNS compared to WT and their BNS-grown counterparts (Fig. 4b,c). A modest reduction in seed set was additionally observed in *glcat14a/b/d-1* relative to WT in BNS (Fig. 4c). Notably, anthers of *glcat14a/b/d* mutants grown in LCNS released almost no pollen, hindering reciprocal crosses to further investigate the reduced seed set.

**Fig 4.**
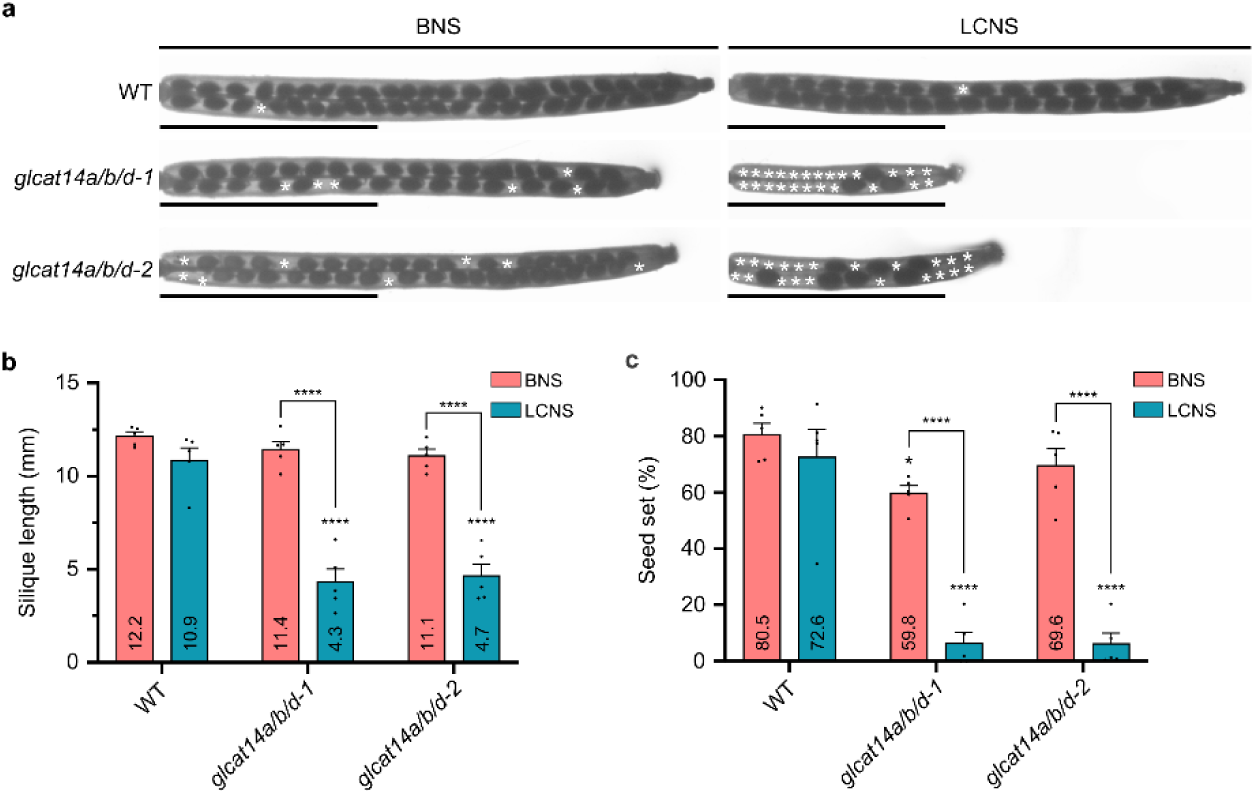
Reduced seed set in *glcat14a/b/d* mutants under low-calcium conditions. **a**, Cleared siliques from seven-week-old wild-type (WT), *glcat14a/b/d-1*, and *glcat14a/b/d-2* plants grown in basal nutrient solution (BNS) and low calcium nutrient solution (LCNS). Areas without seeds are marked with white asterisks. Scale bars, 5 mm. **b**,**c**, Silique length (**b**) and seed set percentage (**c**) of seven-week-old WT and *glcat14a/b/d* mutants under BNS and LCNS. Data are presented as mean + SEM (*n* = 5 plants, 10 siliques per plant). Biological replicates are shown as black dots. Statistical significance was determined by two-way ANOVA followed by Tukey’s test. Asterisks represent statistically significant differences from WT grown in BNS or LCNS, unless otherwise specified (*, *p*<0.0332; ****, *p*<0.0001).

### Low Ca^2+^ exacerbates pollen and pollen tube growth defects in *glcat14a/b/d* mutants

To further assess how Ca^2+^ availability influences male fertility, we examined pollen viability in *glcat14a/b/d* mutants grown under BNS and LCNS. At flower stages 12–13, mutants grown in BNS contained both viable red-magenta PGs and clusters of greenish aborted PGs, whereas mutants grown in LCNS showed mostly aborted PGs that frequently formed clusters (Fig. 5a). Quantification revealed that aborted PGs represented 38–39% of total PGs in *glcat14a/b/d* mutants under BNS, increasing to 76–78% under LCNS, both significantly higher than WT, which displayed predominantly viable PGs (Fig. 5b). Moreover, both mutants showed a significant increase in aborted PGs under LCNS compared with BNS, indicating that low Ca^2+^ exacerbates defects in pollen viability. DAPI staining confirmed the presence of sperm and vegetative nuclei in viable PGs of WT and *glcat14a/b/d* mutants in both nutrient conditions, whereas aborted PGs in the mutants lacked both nuclei (Fig. 5c). At the tricellular stage, anther locules of *glcat14a/b/d* mutants grown under BNS contained collapsed PGs devoid of cytoplasm and retaining only exine remnants, a phenotype absent in WT. Under LCNS, mutants produced only a small number of viable PGs, which were surrounded by numerous collapsed PGs (Fig. 5d).

**Fig 5.**
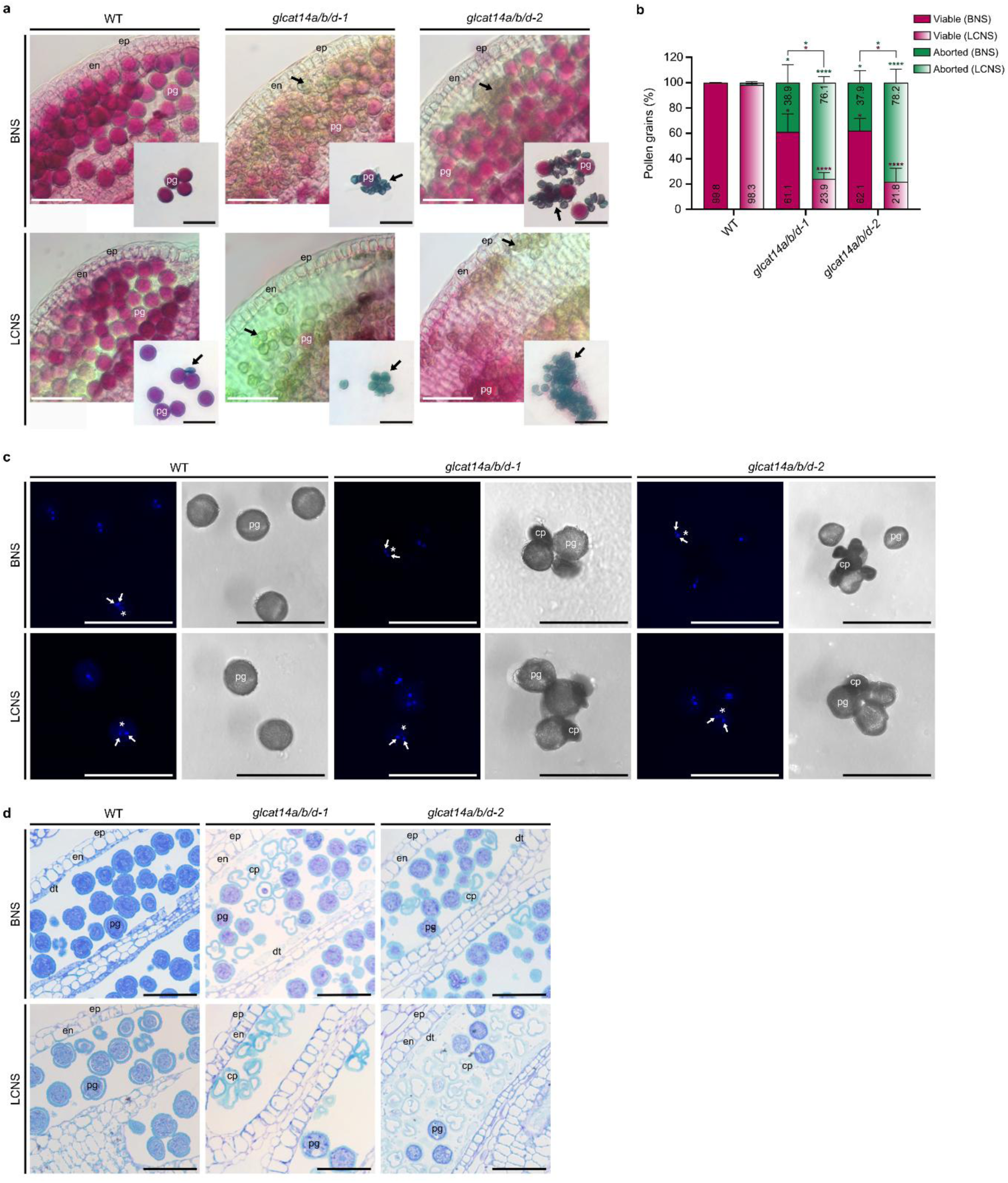
Impaired pollen viability and development in *glcat14a/b/d* mutants under low calcium. **a**, Alexander staining of anthers at flower stage 12 and pollen grains (PGs) of anthers at flower stage 13 of wild-type (WT), *glcat14a/b/d-1*, and *glcat14a/b/d-2* grown with basal nutrient solution (BNS) and low calcium nutrient solution (LCNS). Viable PGs are stained in red-magenta while aborted PGs are stained in blue-green (black arrows). **b**, Viable and aborted PGs percentage of WT and *glcat14a/b/d* mutants grown with BNS and LCNS. Data are presented as mean + SEM (*n* = 3 plants, 500 PGs per plant). Mean values are shown within bars. Statistical significance was determined by two-way ANOVA followed by Tukey’s test. Asterisks represent statistically significant differences from WT grown in BNS or LCNS, unless otherwise specified. Magenta asterisks indicate significance for viable pollen, and green asterisks indicate significance for aborted pollen (*, *p*<0.0332; ****, *p*<0.0001). **c**, DAPI staining of PGs of anthers at flower stage 13 of WT and *glcat14a/b/d* mutants. Sperm cell (white arrows) and vegetative (white asterisks) nuclei are shown in the DAPI channel, alongside the corresponding brightfield channel indicating PG boundaries. **d**, Anther sections at flower stage 12 showing tricellular PG stages of WT and *glcat14a/b/d* plants. Abbreviations: cp, collapsed pollen grain; dt, degenerated tapetum; en, endothecium; ep, epidermis; pg, pollen grain. Images were acquired using a Zeiss Axio Imager.A2 microscope with DIC optics (**a**,**d**) and Leica TCS SP5 II confocal system (**c**). Scale bars, 50 μm.

To determine whether reduced Ca^2+^ availability further impairs pollen performance, we analysed in vitro germination and PT growth of viable PGs from *glcat14a/b/d* mutants grown under BNS and LCNS. Viable PGs of *glcat14a/b/d-1* and *glcat14a/b/d-2* germinated at significantly lower rates than WT under BNS and further reduced under LCNS. Additionally, both mutants showed a significant decrease in germination when grown under LCNS compared with BNS, indicating that low Ca^2+^ further aggravates defects in pollen germination (Extended Data Fig. 5a,b). PTs of mutants were consistently shorter than WT under both BNS and LCNS (Extended Data Fig. 5a,c). In *glcat14a/b/d* mutants, only 24% of total PGs were viable, of which 20% were able to germinate. From these PGs, only 5% produced PTs, which were also substantially shorter than WT. These results indicate that low Ca^2+^ conditions strongly exacerbate defects in both pollen viability and germination, severely limiting mutant fertility.

## Discussion

Male gametophyte development is a critical determinant of plant reproductive success, as pollen viability, germination, and PT growth directly influence fertilisation and seed production^4^. Proper pollen development requires coordination between microspores and the surrounding sporophytic tissues to form structurally competent PGs. AGPs, abundant extracellular glycoproteins present in these tissues^3^, have been implicated in cell wall organisation, signalling, and Ca^2+^ dynamics^9^, yet their precise role in pollen development and fertility remains poorly understood, particularly regarding GlcA residues that mediate Ca^2+^ binding.

Our study demonstrates that glucuronidation of AGPs by GLCAT14A, GLCAT14B, and GLCAT14D is essential for pollen wall formation, viability, and PT growth, and that these processes are sensitive to environmental Ca^2+^ availability. We show that *glcat14a/b/d* mutants have reduced plant growth, shorter siliques with lower seed set, collapsed PGs lacking cytoplasmic content and intine layer, as well as reduced pollen germination and shorter PTs. These defects were exacerbated under low Ca^2+^ conditions and largely attenuated under standard Ca^2+^ levels, highlighting the functional importance of AGP glucuronidation for reproductive success.

We examined the previously reported *glcat14a/b/d* mutants, which contained lower GlcA levels and impaired Ca^2+^ binding^20^. Although these mutants can grow and reproduce normally, indicating that these genes are not essential for survival, they still revealed reproductive defects that phenocopy AGP-deficient mutants. Defective AGP glucuronidation affected overall plant growth, with *glcat14a/b/d* showing delayed development and shorter stems, consistent with previous reports^20^. Similar growth reductions occur in other mutants affecting AGP glycosylation, such as Hyp-O-GALTs, which add the initial galactose residue to AGPs^23^, further supporting the role of AGP glycosylation in plant development.

Silique length and seed set were clearly reduced in *glcat14a/b/d* plants, similar to phenotypes observed in AGP loss-of-function mutants, including AGP6, AGP11^24^, AGP18^25^, AGP19^26^, and FLA3^27^, indicating that impaired AGP function, whether through altered glucuronidation or AGP loss, consistently compromises silique development and reproductive success. Reciprocal crosses showed no significant differences between self- and reciprocal crosses of *glcat14a/b/d* mutants, suggesting that the reduced fertility is not restricted to a single gametophyte and may involve sporophytic contributions.

Male gametophyte defects were evident, with 32% of *glcat14a/b/d* PGs aborting between the uninucleate and bicellular stages, characterised by a shrunken morphology lacking cytoplasmic content and nuclei. A similar phenotype has been reported in AGP mutants, such as *agp6 agp11*^28^, *BRASSINOSTEROID-ASSOCIATED RECEPTOR-LIKE PROTEIN 1* (*BCP1*) antisense line^29^, *FASCICLIN-LIKE AGP 3* (*FLA3*) RNA interference (RNAi) line^27^, and *FLA14* overexpression (OE)^30^ line, which present nearly 50% pollen abortion at the same stages. TEM analysis revealed collapsed PGs in *glcat14a/b/d* lacking the intine layer but retaining an intact exine, whereas viable PGs exhibited irregularities in the intine. Similar intine defects in in *FLA3*-RNAi and *FLA14*-OE lines support a role for these AGPs in intine formation^27,30^. The absence of intine between the uninucleate and bicellular stages leaves microspores structurally unprotected, leading to collapse. A related mechanism involves the nexine layer, whose formation depends on AGPs such as *AGP6*, *AGP11*, *AGP23*, and *AGP40* under the control of TRANSPOSABLE ELEMENT SILENCING VIA AT-HOOK (TEK), with *TEK* expression in the tapetum regulated by the transcription factor ABORTED MICROSPORES (AMS). In *tek* mutants, the absence of nexine disrupts intine deposition and causes microspore collapse^31^, whereas expression of *AGP6* under the *TEK* promoter partially rescues nexine formation^32^. Mutants of *KAONASHI 4* (*KNS4*), which encodes an AGP GALT, further demonstrate that defects in primexine and exine structure correlate with reduced AGP content in developing microspores^33^. These similarities suggest that AGP glycosylation is crucial for pollen wall integrity and that functional AGPs in these layers are essential for PG development.

Collapse PGs in *glcat14abd* mutants also lack cellulose in the intine, similar to aborted PGs in *FLA3*-RNAi and *FLA14*-OE lines^27,30^, as well as in *Brassica campestris Male Fertility 8* (*BcMF8*) and *BcMF18* double antisense mutants, orthologues of *AGP11* and *AGP6*^34^. Supporting a role for AGPs in cellulose deposition, FLA11 and FLA12 influence cellulose deposition^35^, FLA16 regulates cellulose levels^36^, and loss of *FLA4/SALT OVERLY SENSITIVE 5* (*SOS5*) disrupts cellulose patterning in seed mucilage^37^. In *Gossypium hirsutum*, *GhAGP4*-RNAi lines show impaired cellulose deposition during secondary wall synthesis^38^, and *GhFLA1* overexpression or knockdown respectively up- or downregulates *CELLULOSE SYNTHASE 1* (*GhCESA1*), linking AGPs to cellulose biosynthesis^39^. β-Yariv treatment, which binds and disrupts AGPs, also prevents cellulose deposition^40^. *NOVEL MICROGAMETOPHYTE DEFECTIVE MUTANT 1* (*NMDM1*) further links AGP regulation to intine formation and cellulose deposition, as NMDM1 is required for normal expression of *AGP6, AGP23, AGP41, FLA3,* and *FLA12,* all downregulated in *nmdm1*, which lacks cellulose in the intine of aborted PGs^41^. AGPs are proposed as cell wall cross-linkers^9^ via interactions with pectin and hemicellulose, as in the ARABINOXYLAN PECTIN ARABINOGALACTAN PROTEIN 1 (APAP1) complex^42^. Together, these observations support the hypothesis that AGPs contribute to intine formation via cellulose deposition. While a direct molecular link between AGPs and cellulose synthesis remains unproven, recent reviews^9,43^ propose mechanisms by which AGPs may influence cellulose deposition via Ca^2+^-dependent signalling, interactions with pectin–hemicellulose networks, or modulation of cellulose synthase localisation.

In vitro pollen germination and PT growth were reduced in *glcat14a/b/d* mutants, resembling defects reported in *FLA3*-RNAi^27^ and *AGP6 AGP11* double-RNAi lines^24^, respectively, indicating that AGP function is critical for pollen performance. AGP immunolabelling, observed in the same anther tissues reported previously^44^, was slightly reduced in *glcat14a/b/d* mutants, particularly in the epidermis and endothecium. Although some AGP epitopes are partially altered in *glcat14a/b/d* mutants, most AGPs are retained in anther locules, suggesting that reproductive defects may arise from functional impairments, such as altered AGP–Ca^2+^ interactions, rather than from a loss of AGPs. *GLCAT14A/B/D* genes are expressed in sporophytic tissues surrounding the microspores, including the endothecium and epidermis, but not in the gametophyte (unpublished data). We hypothesize that the absence of intine is an indirect sporophytic effect of impaired AGP glucuronidation in sporophytic tissues, which compromises microspore support and leads to collapsed PGs.

Mutants of *GLCAT14A/B/D* exhibit reproductive defects that are highly sensitive to Ca^2+^ availability, exacerbating previous phenotypes under low Ca^2+^ conditions. Restoring Ca^2+^ levels largely suppressed these defects, suggesting that impaired AGP–Ca^2+^ interactions are a primary cause of the defects. This hypersensitivity to Ca^2+^ deprivation is consistent with previous observations in *GLCAT* mutants affecting inflorescence stem growth, trichome branching, and hypocotyl elongation^20^, reinforcing that AGP-mediated Ca^2+^ homeostasis is essential across multiple developmental contexts, consistent with the broad expression and functional diversity of AGPs. Previous work demonstrated that increasing external Ca^2+^ does not rescue phenotypes of pectin-defective mutants, suggesting that the conditional defects in *glcat14a/b/d* are not attributable to altered pectin-mediated Ca^2+^ signalling^20^. Instead, these findings support a direct requirement for AGP glucuronidation in maintaining Ca^2+^ availability for proper pollen wall formation and function. The Ca^2+^ hypersensitivity of *glcat14a/b/d* also contrasts with mutants of Ca^2+^ transporters, such as CYCLIC NUCLEOTIDE-GATED CHANNEL2 (CNGC2) and CATION EXCHANGER1 (CAX1) and CAX3, which tolerate low Ca^2+^ but become dwarfed under high Ca^2+^ due to apoplastic Ca^2+^ overaccumulation^45^. This divergence implies that *glcat14a/b/d* defects reflect impaired AGP-dependent Ca^2+^ buffering rather than altered Ca^2+^ transport. Although Ca^2+^ sensitivity has not been explored in other GTs or AGP mutants, some AGP-deficient lines show phenotypes reminiscent of Ca^2+^ deficiency. For example, *AGP6 AGP11*-RNAi anthers release almost no pollen unless mechanically manipulated^24^, similar to *glcat14a/b/d* under LCNS. Moreover, the methyl-glucuronosyl oligosaccharide AMOR, containing the disaccharide 4-Me-GlcA-β-(1-6)-Gal, induces PT competency to respond to ovular LURE peptides in *Torenia fournieri*^46^, highlighting a role for AGP-derived GlcA in Ca^2+^-mediated signalling^47^. Consistently, Ca^2+^- and signalling-related genes are misregulated in *agp6 agp11* pollen, including upregulation of the *CALMODULIN-LIKE PROTEIN 42* (*CML42*)^48^, supporting a connection between AGPs, Ca^2+^ buffering, and downstream Ca^2+^-dependent pathways in male gametophyte development.

AGPs are modified with GlcA, which binds Ca^2+^ in a reversible and pH-dependent manner, and most are predicted to be anchored to the plasma membrane via a GPI anchor, thus serving as extracellular Ca^2+^ capacitors that generate a layer of Ca^2+^ available for signalling processes^10,14^. Previously, we proposed a model for AGP-mediated Ca^2+^ regulation during PT growth in the transmitting tract^9^, based on previous models^14,47^. In this model, proton efflux via plasma membrane H^+^-ATPases acidifies the apoplast, triggering Ca^2+^ release from PT tip- and transmitting tract-localised AGPs. Released Ca^2+^ enters the cytosol to establish a tip-focused gradient, guiding PT growth and exocytosis, while PT-mediated acidification of the transmitting tract generates a Ca^2+^ gradient that guides PTs toward the ovules. The discharged AGP–Ca^2+^ capacitor is subsequently recharged by recycled Ca^2+^. Extending this concept, Fig. 6 illustrates a generalised AGP–Ca^2+^ capacitor model integrating our current findings. In WT plants under normal Ca^2+^ conditions, local apoplastic acidification releases Ca^2+^ from AGPs, supporting cytosolic Ca^2+^ signalling required for pollen development and PT growth. In *glcat14a/b/d* mutants, reduced AGP glucuronidation lowers Ca^2+^ binding capacity. While normal Ca^2+^ levels partially compensate for this defect, under low Ca^2+^ conditions AGP–Ca^2+^ loading is severely limited, restricting Ca^2+^ signalling and disrupting pollen development and PT growth.

**Fig 6.**
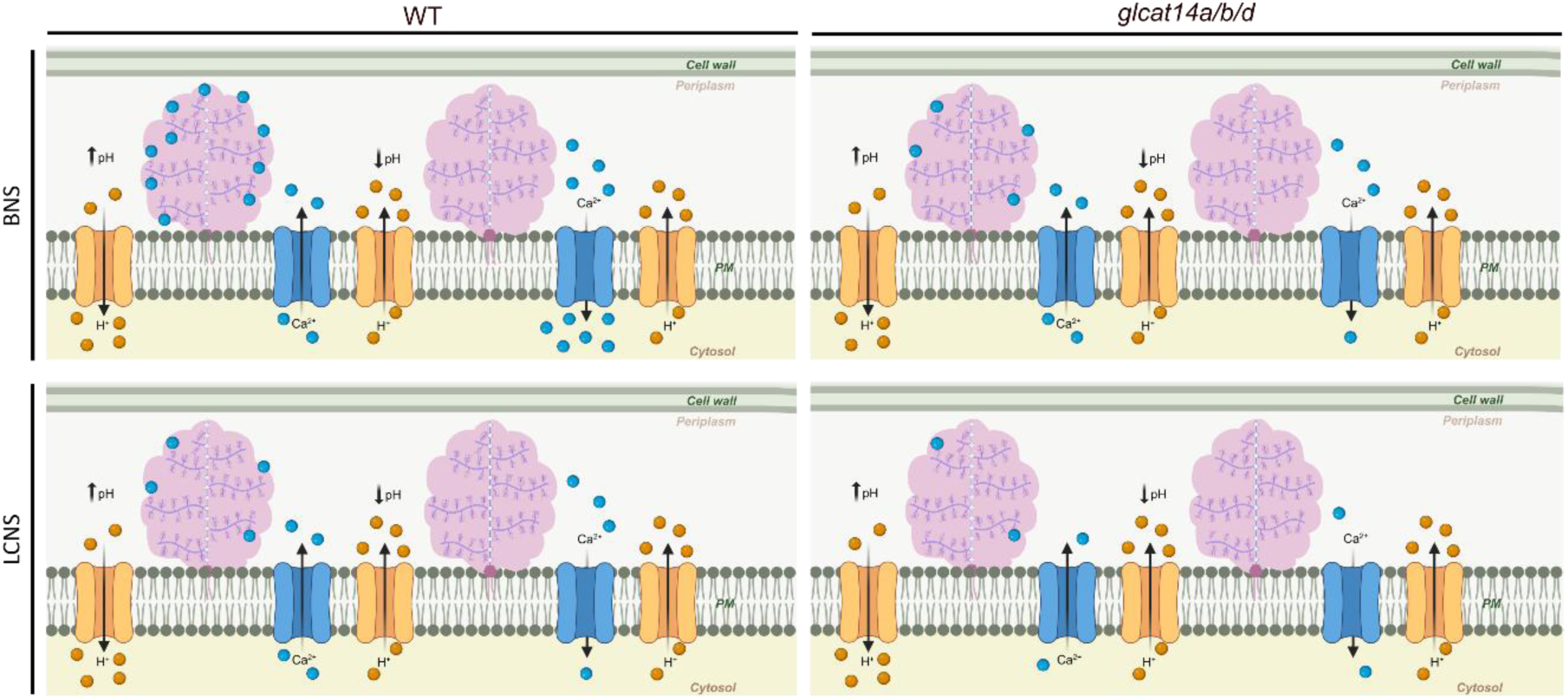
Model of the AGP-Ca^2+^ capacitor in wild-type and *glcat14a/b/d* mutants under low and normal calcium conditions. In wild-type (WT) plants grown with basal nutrient solution (BNS), H^+^ (orange circles) efflux driven by plasma membrane H⁺-ATPases (orange channel) acidifies the apoplast (⭣pH). Under this transiently low periplasmic pH, calcium (Ca^2+^; blue circles) is dissociated from AGPs (pink cotton candy shape), resulting in Ca^2+^ influx into the cytosol through Ca^2+^ channels (blue channels). The resulting rise in cytosolic Ca^2+^ levels drives Ca^2+^-dependent processes. As the pH increases (⭡pH) due to H^+^ influx, AGPs rebind Ca^2+^, restoring the AGP-Ca^2+^ capacitor. In *glcat14a/b/d* mutants grown with BNS, reduced glucuronidation of AGPs lowers their Ca²⁺-binding capacity, limiting the amount of Ca^2+^ that can be released from GlcA-deficient AGPs. This defect is partially compensated by the higher external Ca^2+^ available in BNS, allowing some Ca²⁺-dependent processes to proceed. In WT plants grown with low calcium nutrient solution (LCNS), low external Ca^2+^ restricts the number of Ca^2+^ ions available for AGP binding. Nevertheless, the Ca^2+^ released from AGPs still reaches the minimum cytosolic levels required to sustain Ca^2+^-dependent signalling. In *glcat14a/b/d* mutants grown in LCNS, both impaired AGP glucuronidation and low external Ca^2+^ further reduce AGP–Ca^2+^ loading. Consequently, the Ca²⁺ released from AGPs is insufficient to properly drive Ca²⁺-dependent processes, leading to severe developmental and reproductive defects. Created with BioRender.com.

This study reveals the functional importance of GlcA residues in AGPs, which modulate Ca^2+^ signalling and support cellulose deposition and pollen wall integrity. Furthermore, glucuronidation of AGPs by GLCAT14A/B/D, and the resulting Ca^2+^ binding, is essential for male gametophyte viability, PT growth, and successful fertilisation, highlighting AGPs as dynamic extracellular Ca^2+^ reservoirs in plant reproduction. Beyond the male gametophyte, these results suggest a broader principle whereby AGPs can serve as regulated Ca^2+^ stores to coordinate developmental processes. Future work should investigate the prevalence of AGP–Ca^2+^ capacitors across tissues and species, explore their integration with other Ca^2+^-dependent signalling pathways in plant growth and reproduction, and determine whether the remaining eight members of the GT14 family contribute to AGP-mediated Ca^2+^ regulation.

## Methods

### Plant material and growth conditions

All *Arabidopsis thaliana* (L.) Heynh. plants used in this study were in the Col-0 ecotype. WT seeds were obtained from Eurasian Arabidopsis Stock Centre (NASC). The T-DNA mutant lines *glcat14a glcat14b glcat14d-1* (*glcat14a/b/d-1*) and *glcat14a glcat14b glcat14d-2* (*glcat14a/b/d-2*) were previously reported^20^. Triple mutants were generated by crossing *glcat14d-1* (GK-363F05.01, N377379) or *glcat14d-2* (GK-508D01.01, N326773) T-DNA lines with *glcat14a glcat14b*, which was obtained by crossing *glcat14a-2* (SALK_043905, N543905) with *glcat14b-1* (SALK_080923, N580923)^20^. Seeds were stratified for 48 h at 4 °C in the dark and sown directly on soil (COMPO SANA). Plants were grown under long-day conditions (16 h light at 22 °C and 8 h darkness at 18 °C), with 50–60% relative humidity and 180 µmol m⁻² s⁻¹ light intensity. Mutant lines were screened by PCR genotyping and RT–qPCR. Seeds were collected and propagated for subsequent analyses.

Hydroponic growth conditions followed a previously described protocol^20^. Seeds were surface-sterilized in 80% (v/v) ethanol containing 1% (v/v) NaOCl and 1% (v/v) Tween-20, followed by sequential washes in 65%, 80% and 100% (v/v) ethanol. Sterilised seeds were sown onto half-strength MS medium with vitamins [0.44% (w/v); Duchefa Biochemie], supplemented with 0.1% (w/v) MES and 1% (w/v) sucrose, adjusted to pH 5.8, and solidified with 0.9% (w/v) plant agar. Plates were cold-stratified for 48 h at 4 °C in darkness and then placed vertically (slightly inclined) under long-day conditions for two weeks. New squared pots with drainage holes were filled with a 1 cm layer of vermiculite and a rockwool cube (Cultilene). Drainage holes were blocked with soft foam to retain vermiculite while allowing nutrient solution flow. Pots were washed with deionised water for 2 days, with daily water changes, and covered with aluminium foil to prevent evaporation and contamination. Two-week-old seedlings were transferred through openings in the foil, with four plants per pot. Each pot was watered with 250 mL of nutrient solution containing either 0.2 mM Ca^2+^ (low calcium nutrient solution, LCNS) or 2 mM Ca^2+^ (basal nutrient solution, BNS) every fifth day (Supplementary Table 3). Plants were grown under long-day conditions (16 h light at 22 °C and 8 h darkness at 18 °C).

### Genotyping

To confirm T-DNA insertions and identify homozygous mutants, genomic DNA was extracted from rosette leaves^49^, and PCR-based genotyping was performed using DreamTaq DNA Polymerase (Thermo Fisher Scientific) according to the manufacturer’s instructions, with gene-specific primers for *GLCAT14A*, *GLCAT14B*, and *GLCAT14D*, together with the corresponding T-DNA left-border primers (Supplementary Table 4). T-DNA insertion sites were further confirmed by DNA sequencing. The *glcat14a/b/d-1* and *glcat14a/b/d-2* triple mutants carried two T-DNA insertions in opposite orientations in the 2^nd^ exon of *GLCAT14A* and in the 4^th^ exon of *GLCAT14B*. In *GLCAT14D*, *glcat14a/b/d-1* contained a single insertion in the 3^rd^ intron, whereas *glcat14a/b/d-2* harboured two insertions in opposite orientations, one in the 3^rd^ intron and one in the 4^th^ exon (Extended Data Fig. 1a).

### RNA extraction, cDNA synthesis, and qPCR analysis

Inflorescences from five-week-old plants, comprising flowers at developmental stages 1–15^21,22^ were collected from three individual plants per genotype (*n* = 3), immediately frozen in liquid nitrogen, and stored at −80 °C for subsequent experiments. Total RNA was extracted using PureZol RNA Isolation Reagent (Bio-Rad) following the manufacturer’s protocol. One microgram of RNA was treated with DNase I, RNase-free (Thermo Fisher Scientific) according to the manufacturer’s instructions. cDNA was synthesised using the RevertAid First Strand cDNA Synthesis Kit (Thermo Fisher Scientific) with oligo(dT)18 primers, following the manufacturer’s guidelines, and diluted to 5 ng/μL in nuclease-free water (Sigma-Aldrich) for quantitative real-time PCR (qPCR) analysis.

qPCR primers for *GLCAT14A*, *GLCAT14B*, and *GLCAT14D* were designed (Supplemental Table 5) and validated as described previously^50^. Primer efficiency, correlation coefficient (R^2^), and melting temperature were determined from a standard curve generated using a three-point tenfold dilution series of cDNA from WT inflorescences (Supplementary Table 6). qPCR reactions were performed as reported previously^50^ using 2× SsoAdvanced Universal SYBR® Green Supermix (Bio-Rad) with three technical replicates per biological replicate and non-template controls on a CFX96 Touch Real-Time PCR Detection System (Bio-Rad). Quantitative cycle (Cq), baseline correction, and threshold were automatically calculated using CFX Maestro 2.0 (Bio-Rad). Target gene expression was normalised to three validated reference genes (*HIS3.3*, *ACT7*, and *YLS8*)^50^ with stability values (*M*) of 0.16, 0.16, and 0.20, respectively, and normalised relative gene expression was calculated using the 2^−ΔΔCt^method^51^.

### Plant and floral phenotyping

Flowering time was recorded as days to anthesis, defined as the stage when the first anther dehisces to release pollen. Twelve plants (*n* = 12) per genotype were analysed. Main inflorescence stem, stamen, and pistil lengths were measured using FIJI^52^. Stem length was measured in twelve plants (*n* = 12) per genotype. For each genotype and developmental stage, five flowers from three plants (*n* = 3) were analysed, with 5 pistils and 24–31 stamens measured per plant. Plants and stems were captured using a Nikon D3300 digital camera. Flowers from stages 12–15 were imaged using a Zeiss Stemi 305 stereomicroscope with an Axiocam 208 color camera (1.0× camera adapter, 0.75× objective, 2.0× optovar) and ZEN software (exposure = 56 ms).

### Seed set analysis

Siliques from 7-week-old plants were collected for silique length measurements, seed set analysis, and clearing. For each plant, siliques were harvested from the main stem after discarding the five oldest and five youngest siliques. Silique length was measured under a Nikon SMZ800 stereo microscope using millimetre paper, with an uncertainty of 0.5 mm. For each genotype, five siliques (technical replicates) from five plants (*n* = 5) were analysed. Siliques were dissected with hypodermic needles, and seeds were counted and classified as fully developed, white (arrested at the globular stage), aborted (desiccated before white seeds), or aborted ovules. Seed set was calculated as the ratio of mature seeds to total seeds. Ten siliques (technical replicates) from five plants (*n* = 5) of each genotype were analysed.

For self- and reciprocal crosses, closed flower buds at flower stage 12 were emasculated and, after 24 h, hand-pollinated using fresh pollen from open flowers at stage 13–14. Siliques were collected ten days after pollination (DAP), and silique length and seed set were determined as described above. Five siliques (technical replicates) from three plants (*n* = 3) per cross were analysed.

Siliques were fixed overnight at 4 °C in 1:9 (v/v) glacial acetic acid (Fisher Scientific) and 100% (v/v) ethanol (Fisher Scientific), then dehydrated in 90% (v/v) ethanol for 10 min and 70% (v/v) ethanol for 10 min. Cleared siliques were imaged at 0.8× magnification using a Nikon SMZ800 stereo microscope equipped with a Nikon DXm1200F digital camera and Nikon ACT-1 software (exposure = 300 ms).

### Fluorescent immunolocalisation

Closed flower at stages 9–12 were fixed, dehydrated, and embedded in LR White resin following the protocol previously described^53^. Briefly, samples were fixed in a formaldehyde–glutaraldehyde–PIPES solution, dehydrated through a graded ethanol series, and embedded in LR White resin (Agar Scientific) through a crescent series of ethanol:resin ratios. Resin polymerisation was performed in gelatine capsules (Agar Scientific) for 48 h at 60 ° C. Anther thin sections (500 nm) were cut using a Leica EM UC7 Ultramicrotome with a Diatome diamond knife (Agar Scientific) and mounted on diagnostic microscope slides with 8 wells (Assistant). Flowers from three plants (*n* = 3) of each genotype were analysed. Fluorescent immunolocalisation was performed as previously described^53^. In brief, each well was incubated with 25 μL of filtered blocking solution [5% (w/v) skim milk in 1× PBS] for 10 min and washed twice with PBS (10 min each). Wells were then incubated with 25 μL of primary antibody (1:5) for 2 h at room temperature in a humid chamber, followed by overnight incubation at 4 °C. Negative controls were treated with blocking solution only. After washing twice with PBS and once with deionized water (10 min each), wells were incubated with 25 μL of FITC-conjugated anti-rat IgG secondary antibody (1:100; Sigma-Aldrich) for 4 h in the dark at room temperature in a humid chamber. Wells were washed as above, then stained with a drop of 0.01% (w/v) filtered calcofluor white (Sigma-Aldrich) and mounted with a drop of Vectashield (Vector Laboratories). Primary antibodies included anti-AGP monoclonals JIM8, JIM13, LM2, and LM14, and anti-pectin monoclonals JIM5 and JIM7 (Plant Probes). JIM13^54,55^ and JIM8^54,56^ detect AGP glycans, LM2^55,57,58^ detect β-1,6-galactan with terminal GlcA, LM14^56,59^ detect GlcA in AGP glycan, JIM5^60–63^ recognises partially methyl-esterified and un-esterified homogalacturonan, and JIM7^61–63^ recognises partially methyl-esterified homogalacturonan.

### Pollen viability and content analysis

Pollen viability was assessed using Alexander staining^64–66^ following the published protocols, with minor modifications. Closed flowers at stage 12 were fixed overnight at 4 °C in Carnoy’s fixative^65^, dehydrated in 70% (v/v) ethanol, and dissected using hypodermic needles. Stamens from closed flowers and PGs from freshly opened flowers at stage 13 were mounted on glass microscope slides in Alexander’s stain, incubated at 45 °C for 15 min in the dark, and covered with a coverslip for observation. Five closed flowers and one opened flower containing 500 PGs (technical replicates) from three plants (*n* = 3) of each genotype were analysed. For toluidine blue staining, anther sections (1 µm) were mounted on glass microscope slides, stained with 1% (w/v) toluidine blue (Agar Scientific) at 50 °C, and mounted in DPX (Sigma-Aldrich).

Pollen content was assessed by DAPI staining^66^, with the following modifications. Pollen from freshly opened flowers was released onto a glass microscope slide, and 1 drop of Fluoroshield containing DAPI (Sigma-Aldrich) was added. Samples were incubated in the dark for 5 min before placing a coverslip. One opened flower (technical replicate) of three plants (*n* = 3) of each genotype was analysed.

For pollen wall staining, wells with thin anther sections (500 nm) were washed three times with 1× PBS (5 min each) and incubated with either 0.1% (w/v) DABS^66^ or 1 mg/mL GFP-CBM3 (Carbohydrate Binding Module 3C from *Clostridium thermocellum*, NZYtech) for 1 h at room temperature. For calcofluor white staining, a drop of 0.01% (w/v) filtered calcofluor was applied and slides were mounted in Vectashield.

### Pollen in vitro germination

Pollen germination assays were performed on solid pollen germination medium (PGM), prepared by supplementing liquid PGM^67,68^ with 1.5% (w/v) low-melting agarose (Fisher Scientific). Three 20 × 15 mm rectangles were drawn on a glass microscope slide using an A-PAP pen (Daido Sangyo), and 500 µL of solid PGM was spread within each rectangle to form a flat layer^69^. PGs from three freshly opened flowers at stage 13 were released onto the medium, one flower per rectangle. Slides were placed in a humid chamber and incubated in the dark at 22 °C for 6 h to allow pollen germination and PT growth. Three flowers (technical replicates) from three plants (*n* = 3) per genotype were analysed. For each plant, 150 PGs (total 450 per genotype) were scored as germinated if a PT emerged. Germination percentage was calculated as the ratio of germinated to total PGs. Collapsed PGs, identified by their shrunken morphology, were excluded from scoring. PT length was measured for 50 PTs per plant (total 150 per genotype) using FIJI.

### Transmission electron microscopy (TEM)

Pistils and stamens at flower stage 12 were fixed in fixative solution [2% (v/v) formaldehyde, 2% (v/v) glutaraldehyde fixative in 1× PHEM buffer [60 mM PIPES buffer (Sigma-Aldrich), 25 mM HEPES (Fisher Bioreagents), 10 mM EDTA (Sigma-Aldrich), 2 mM MgCl_2_ (BDH Chemicals) adjusted to pH 7.2 with NaOH (ITW Reagents), and 0.001% (v/v) Tween 20 (VWR)], vacuum-infiltrated for 20 min (−70 kPa), and fixed for 2 h 40 min at room temperature, followed by overnight fixation at 4 °C. Samples were washed with 1× PHEM buffer (five times, 10 min) and post-fixed in 2% (v/v) osmium tetroxide overnight at 4 °C. After washing with deionized water (five times, 10 min), samples were dehydrated in a graded ethanol series (30%, 50%, 70%, 80%, and 90%; 15 min each), followed by three washes in 100% ethanol (10 min each). Samples were washed twice 100% (v/v) acetone (Fisher Scientific; 20 min) and embedded in increasing concentrations of epoxy resin [48% (v/v) Agar 100 resin, 52% (v/v) MNA, 2% (v/v) DMP-30 (Agar Scientific)] in acetone (1:3, 1:1, 3:1, and 1:0; 12 h each at room temperature). Samples were further incubated twice in pure resin (12 h each) and resin was polymerised for 24 h at 60 °C. Ultrathin sections (60—80 nm) were cut on a Leica EM UC7 Ultramicrotome, mounted on mesh copper grids, stained with uranyl acetate substitute and lead citrate (Electron Microscopy Sciences; 5 min each). Sections were examined in a JEOL JEM 1400 transmission electron microscope equipped with STEM detector and EDS system (120 kV), and images were acquired using an Orius CCD digital camera (Gatan). Three closed flowers (technical replicates) from three plants (*n* = 3) of each genotype were analysed.

### Microscopy

DIC images were acquired using a Zeiss Axio Imager.A2 upright microscope with DIC optics, equipped with a Zeiss AxioCam MRc 3 camera and a 60N-C 2/3 0.63× camera adapter. Plan-Neofluar 20×/0.5 NA (for PTs) or 40×/0.75 NA objectives (for anthers and PGs) were used. Images were acquired with Zeiss Zen 2012 software.

Epifluorescence images were captured using a Leica DMLB upright epifluorescence microscope equipped with a Nikon DS-Ri2 color camera and an N PLAN 40×/0.65 objective. A UV-2A filter block (excitation 330–380 nm) was used to detect DABS. All images within a single experiment were acquired with identical gain (2.8×) and exposure (200 ms) settings. Images were captured using Nikon NIS-Elements BR 4.60.00 software.

Confocal images were acquired using Leica TCS SP5 II and SP8 systems equipped with an argon laser. For the SP5 II, fluorescent signals were acquired using an HC PL APO CS 10×/0.40 NA DRY UV objective (for pistils) or a 63×/1.30 NA CORR GLYC objective (for PGs and ovules). Laser power was 10–14%, with the 405 nm line at 20% for DABS and 60% for DAPI. Emission was collected at 417–496 nm (PMT1, 0% offset, HV: 715 V for pistils, 851 V for ovules, and 956 V for PGs). For the SP8, fluorescent signals were acquired using an HC PL APO CS2 10×/0.40 DRY objective (for pistils) or a Fluotar VISIR 25×/0.95 WATER objective (for anthers). Laser power was 9.5%, with the 405 nm line at 2% (for pistils) or at 29.5% and the 488 nm line at 4.7% (for anthers). Emission was collected at 415–478 nm (PMT1, 0% offset, gain 650 for anthers and 700 for pistils) for DABS and calcofluor, and at 498–580 nm (PMT2, 0% offset, gain 700) for GFP–CBM3. The confocal pinhole was 1 AU, scan speed 400 Hz, line average 4, and images were captured at 1024 × 1024 pixels. All confocal images were acquired using Leica LAS AF software.

### Statistical analysis

Statistical analyses and graph generation were performed using GraphPad Prism 10. Data normality was assessed using the Shapiro–Wilk test, and homogeneity of variances was evaluated using the Brown–Forsythe test prior to performing any ANOVA. Depending on data distribution, parametric or non-parametric tests were applied, with significance set at α = 0.05. The specific tests used for each dataset are indicated in the corresponding figure legends. For data showing normal distribution and homogeneous variances, statistical significance was determined by one-way or two-way ANOVA followed by Dunnett’s or Tukey’s multiple comparisons test, depending on whether comparisons were made only against the control or among all groups. When one group deviated from normality, differences among genotypes were evaluated using the Kruskal–Wallis test followed by Dunn’s multiple comparisons test. For qPCR analysis, expression values were tested for normality using the Shapiro–Wilk test, and statistical differences between genotypes were assessed by one-way ANOVA with Tukey’s multiple comparisons test in CFX Maestro 2.0 (Bio-Rad). Asterisks represent statistically significant differences from WT (*, *p*<0.0332; **, *p*<0.0021; ***, *p*<0.0002; ****, *p*<0.0001).

### Image processing and presentation

Images were processed according to their type. Cleared siliques were converted to grayscale and brightness was adjusted in Adobe Photoshop CC 2019. RGB white balance for Alexander- and toluidine blue-stained sections (DIC images) was corrected using the *white balance correction_1.0* FIJI macro (V. Bindokas, 2006; modified by P. Mascalchi, 2017). In vitro PT DIC images were adjusted for window and level in FIJI. Autofluorescent anther sections and DAPI-stained PG confocal images were background-subtracted using a rolling ball radius of 200 pixels and contrast-enhanced at 0.01% saturation in FIJI. Calcofluor-, DABS-, and GFP–CBM3-stained anther sections were processed similarly, with contrast enhancement at 1% saturation. Immunolocalisation images were processed similarly, with contrast enhancement at 0.1–0.2% saturation. All images were arranged into figure panels in Photoshop and final figure assembly, including scale bars and annotations, was performed in Inkscape 1.2.2.

## Data availability

All data supporting this study are included in the paper and Supplementary Information. Source data are provided.

## Acknowledgments

We acknowledge the support of the i3S Scientific Platforms Advanced Light Microscopy (ALM) and Histology and Electron Microscopy (HEMS). We thank Dr. Federico Lopez-Hernandez for providing *Arabidopsis* mutant seeds and MSc. Sara Foubert-Mendes for assistance with microtome sectioning. This work was supported by the “Contrato-Programa” UID/04050/2025 funded by FCT I.P. https://doi.org/10.54499/UID/04050/2025. J.S. has received funding from “la Caixa” Foundation (ID 100010434), under the agreement LCF/BQ/DR20/11790010. M.J.F.’s research was supported by an FCT PhD grant, SFRH/BD/143579/2019.

## Author contributions

Conceptualization, J.S.; Methodology, J.S.; Validation, J.S.; Formal Analysis, J.S.; Investigation, J.S., M.J.F; Writing – Original Draft, J.S.; Writing – Review & Editing, M.J.F, P.D., M.R.T., M.M.R.C., S.C; Visualization, J.S.; Supervision, M.R.T., M.M.R.C., S.C.; Funding Acquisition, J.S., M.J.F. and S.C.

## Extended Data Figures

**Extended Data Fig 1.**
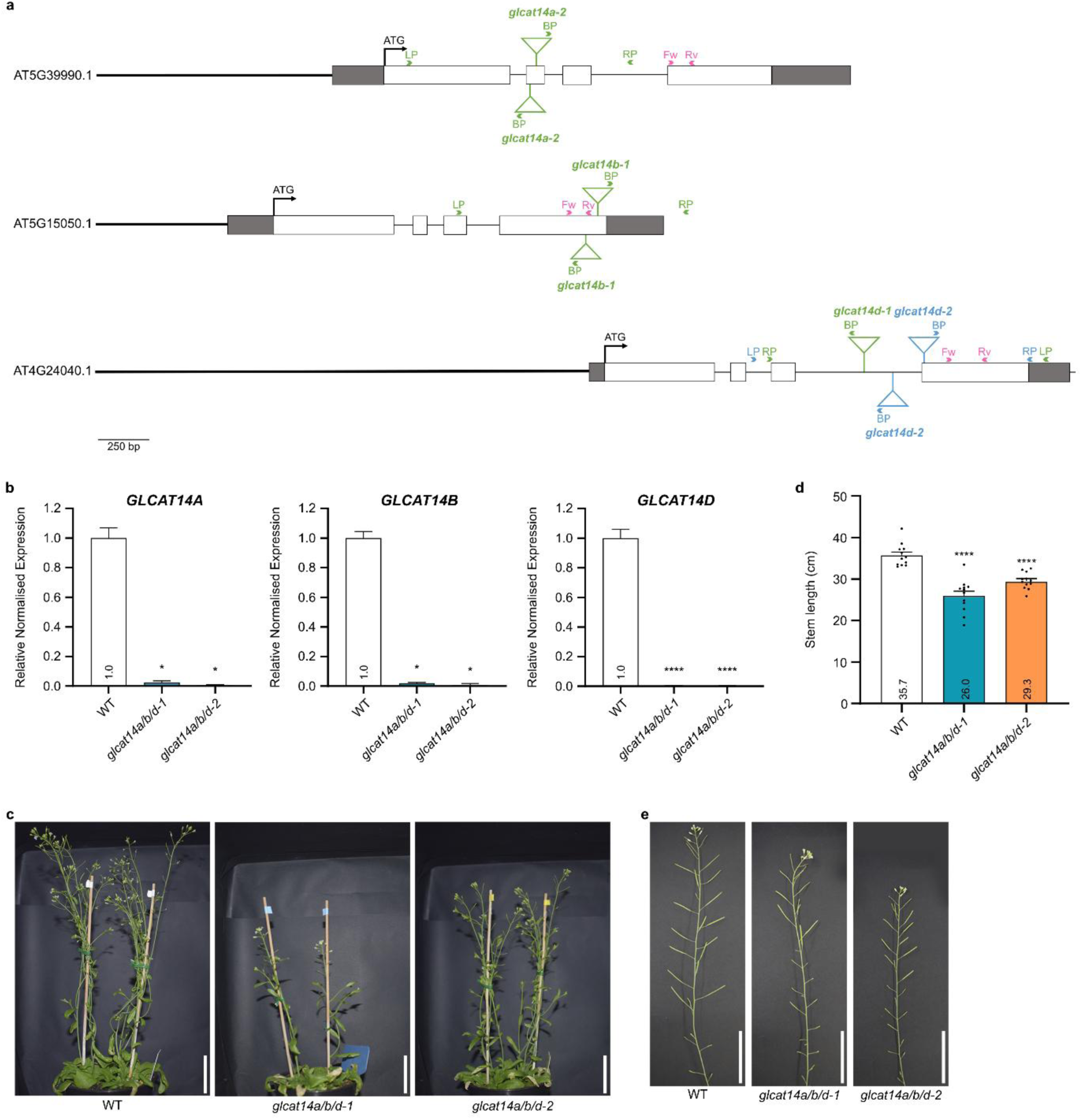
Growth and fertility defects in *glcat14a/b/d* mutants. **a**, Gene structures of *GLCAT14A* (AT5G39990.1), *GLCAT14B* (AT5G15050.1), and *GLCAT14D* (AT4G24040.1) showing T-DNA insertion sites (inverted green and blue triangles) in *glcat14a-2*, *glcat14b-1*, *glcat14d-1*, and *glcat14d-2*. Promoters (thick black lines), untranslated regions (grey boxes), exons (white boxes), and introns (thin lines) are shown. Arrowheads indicate the positions of genotyping (green and blue) and RT-qPCR (pink) primers. **b,** RT-qPCR analysis of *GLCAT14A*, *GLCAT14B*, and *GLCAT14D* transcript levels in inflorescences of wild-type (WT), *glcat14a/b/d-1*, and *glcat14a/b/d-2* plants. Data represent normalised gene expression relative to WT and are shown as mean + SEM (*n* = 3 plants). **c**,**d**, Representative images (**c**) and stem length (**d**) of six-week-old WT and *glcat14a/b/d* plants. **d**, Data are presented as mean + SEM (*n* = 12 plants). Biological replicates are shown as black dots. Statistical significance was determined by one-way ANOVA followed by Tukey’s test (**b**) or Dunnett’s test (**d**). Mean values are shown within bars. Asterisks represent statistically significant differences from WT (**, *p*<0.0021; ****, *p*<0.0001). **e**, Images of seven-week-old stems with siliques WT and *glcat14a/b/d* plants. **c**,**e**, Scale bars, 5 cm.

**Extended Data Fig 2.**
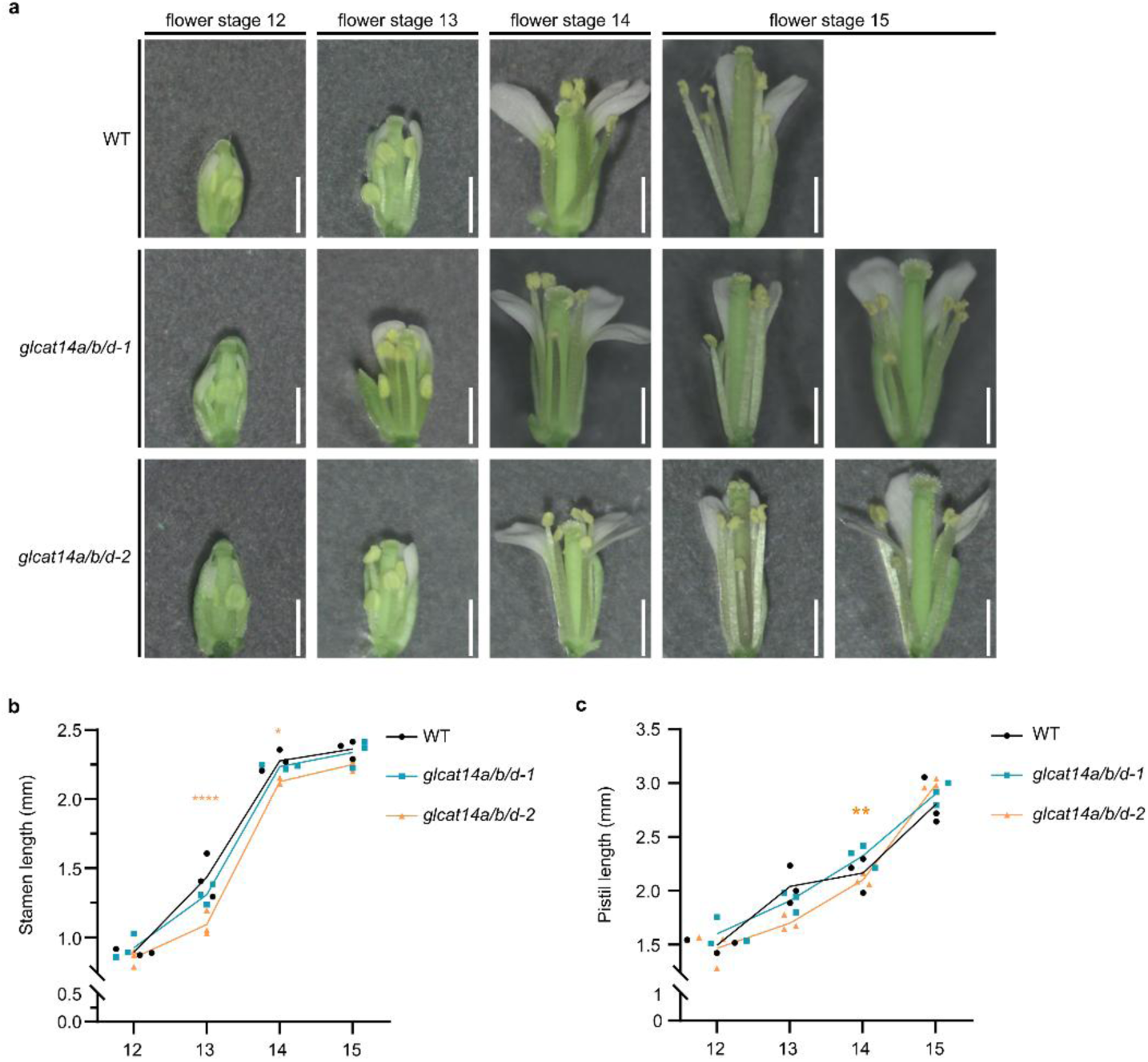
Floral organ morphology and growth in *glcat14a/b/d* mutants. **a**, Representative images of flowers at stages 12–15 of wild-type (WT), *glcat14a/b/d-1*, and *glcat14a/b/d-2* plants. Scale bars, 1 mm. **b**,**c**, Stamen (**b**) and pistil (**c**) length in flowers at stages 12–15 from WT and *glcat14a/b/d* plants. Data are presented as mean values (*n* = 3 plants, 5 pistils per plant for **b** and 24–31 stamens per plant for **c**). Biological replicates are shown as symbols, and lines onnect mean values across stages. Statistical significance was determined by two-way ANOVA followed by Dunnett’s test. Asterisks represent statistically significant differences from WT (*, *p*<0.0332; **, *p*<0.0021; ****, *p*<0.0001).

**Extended Data Fig 3.**
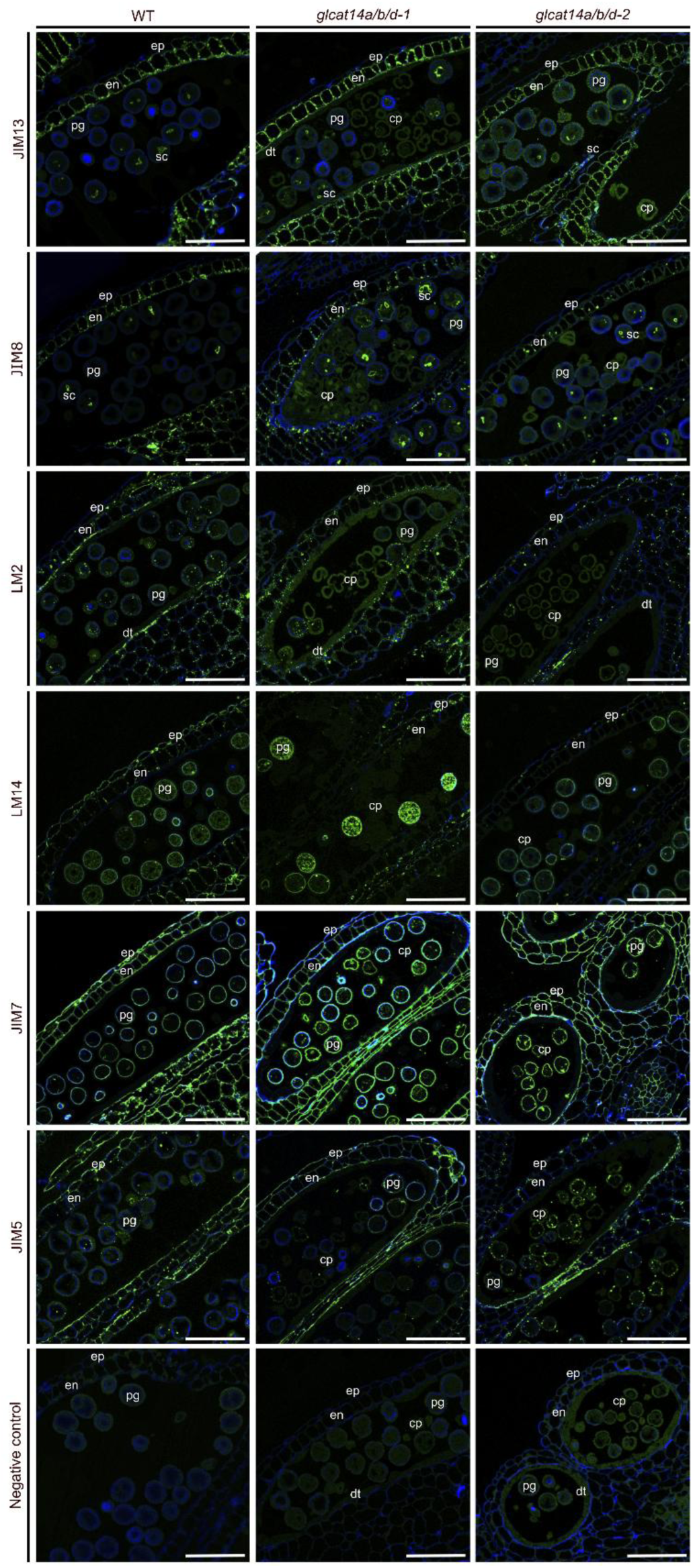
AGP localisation in WT and *glcat14a/b/d* mutants. Immunolocalisation of AGPs using monoclonal antibodies JIM13, JIM8, LM2, and LM14 in anthers sections at flower stage 12 of wild-type (WT), *glcat14a/b/d-1*, and *glcat14a/b/d-2*. A negative control without primary antibody is included. Abbreviations: cp, collapsed pollen grain; dt, degenerated tapetum; en, endothecium; ep, epidermis; pg, pollen grain; sc, sperm cells. Images were acquired using a Leica SP8 confocal systems. Scale bars, 50 μm.

**Extended Data Fig 4.**
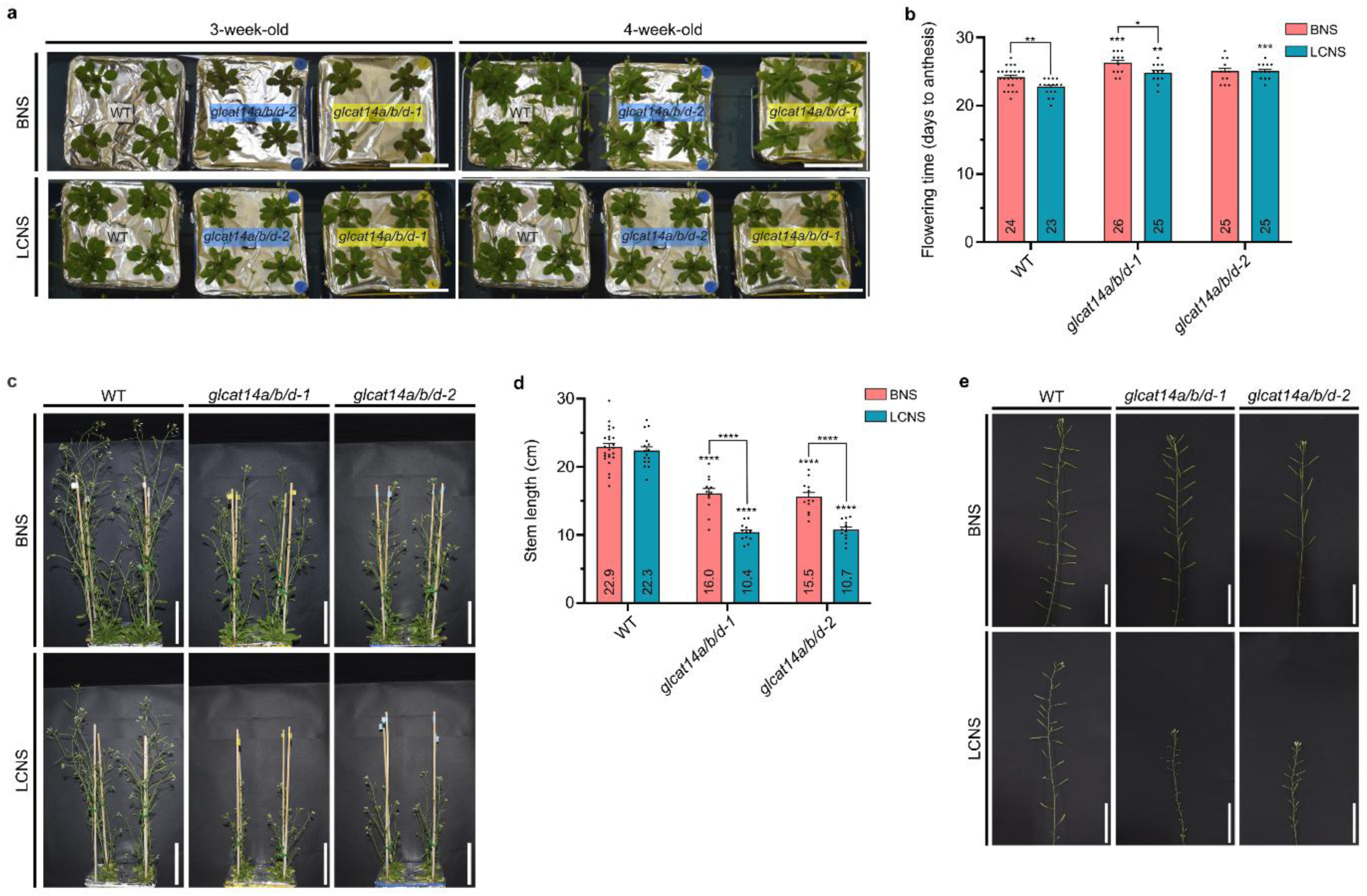
Growth and fertility defects in *glcat14a/b/d* mutants under low-calcium conditions. **a**, Representative images of three- and four-week-old wild-type (WT, white), *glcat14a/b/d-1* (yellow), and *glcat14a/b/d-2* (blue) plants grown in BNS and LCNS. **b**, Flowering time (days to anthesis) of WT and *glcat14a/b/d* mutants under BNS and LCNS. **c**, Images of five-week-old WT and *glcat14a/b/d* plants grown in BNS and LCNS. **d**, Stem length of seven-week-old WT and *glcat14a/b/d* mutants under BNS and LCNS. Data are presented as mean + SEM (*n* = 12 plants, *n* = 24 plants for WT in BNS and *n* = 16 plants in LCNS). Biological replicates are shown as black dots. Statistical significance was determined by two-way ANOVA followed by Tukey’s test. Asterisks represent statistically significant differences from WT grown in BNS or LCNS, unless otherwise specified (*, *p*<0.0332; **, *p*<0.0021; ***, *p*<0.0002; ****, *p*<0.0001). **e**, Images of seven-week-old stems with siliques WT and *glcat14a/b/d* plants grown with BNS and LCNS. Scale bars, 5 cm.

**Extended Data Fig 5.**
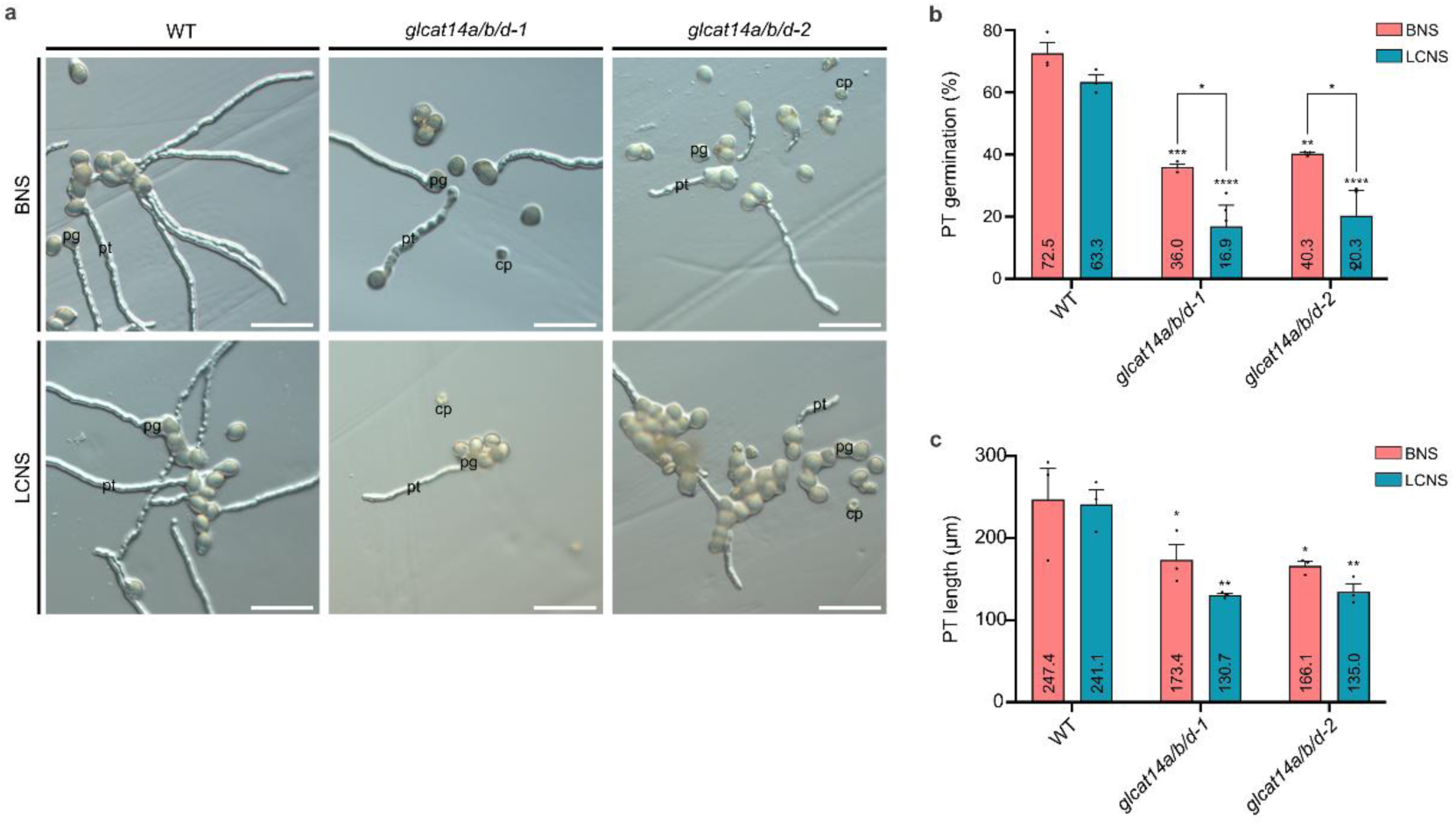
Reduced pollen germination and tube growth in glcat14a/b/d mutants under low calcium. **a**, *In vitro* pollen germination of PGs of wild-type (WT), *glcat14a/b/d-1*, and *glcat14a/b/d-2* grown with BNS and LCNS. Abbreviations: cp, collapsed pollen grain; pg, pollen grain; pt, pollen tube. Images were acquired using a Zeiss Axio Imager.A2 microscope with DIC optics Scale bars, 100 μm. **b**,**c**, Pollen germination percentage (**b**) and pollen tube (PT) length (**c**) of WT and *glcat14a/b/d* mutants *2* grown with BNS and LCNS. Data are presented as mean + SEM [*n* = 3 plants, 150 PGs per plant for (**b**) and 50 PTs per plant for (**c)**]. Biological replicates are shown as black dots. Mean values are shown within bars. Statistical significance was determined by two-way ANOVA followed by Tukey’s test. Asterisks represent statistically significant differences from WT (*, *p*<0.0332; **, *p*<0.0021; ***, *p*<0.0002 ****, *p*<0.0001).

## Supplementary information

**Supplementary Table 1.** Summary of growth and reproductive phenotypes of WT and *glcat14a/b/d* mutants. Values are presented as mean ± SEM. Statistical significance of differences between each mutant and WT is indicated by *p*-values. Abbreviations: ns, not significant.

| Phenotype/Genotype | WT | *glcat14a/b/d-1* | *p* | *glcat14a/b/d-2* | *p* |
| --- | --- | --- | --- | --- | --- |
| RT-qPCR expression (relative to WT) | 1 | 0 | <0.0001 | 0 | <0.0001 |
| Stem length (mm) | 35.7 ± 0.8 | 26.0 ± 1.1 | <0.0001 | 29.3 ± 0.6 | <0.0001 |
| Stamen length (stage 13, mm) | 1.44 ± 0.04 | 1.31 ± 0.03 | ns | 1.09 ± 0.03 | <0.0001 |
| Stamen length (stage 14, mm) | 2.28 ± 0.04 | 2.2 ± 0.04 | ns | 2.13 ± 0.04 | <0.0332 |
| Pistil length (stage 13, mm) | 2.04 ± 0.12 | 1.9 ± 0.06 | ns | 1.70 ± 0.04 | <0.0021 |
| Silique length (mm) | 16.3±0.3 | 13.2 ± 0.5 | <0.0002 | 13.7 ± 0.4 | <0.0002 |
| Seed set (%) | 99.1 ± 0.3 | 85.0 ± 5.7 | <0.0332 | 81.4 ± 5.6 | <0.0021 |
| Viable PGs (%) | 99.6 ± 0.3 | 68.3 ± 1.7 | <0.0001 | 68.1 ± 0.8 | <0.0001 |
| Aborted PGs (%) | 0.4 ± 0.3 | 31.7 ± 1.7 | <0.0001 | 31.9 ± 0.8 | <0.0001 |
| Pollen germination (%) | 63.5 ± 3.3 | 35.1 ± 5.0 | <0.0021 | 35.3 ± 5.0 | <0.0021 |
| PT length (μm) | 290.7 ± 6.9 | 188.7 ± 8.2 | <0.0001 | 190.2 ± 4.5 | <0.0001 |

**Supplementary Table 2.** Summary of growth and reproductive phenotypes of WT and *glcat14a/b/d* mutants grown in BNS and LCNS. Values are presented as mean ± SEM. Statistical significance of differences between each mutant and WT under the same condition is indicated by *p*-values. Abbreviations: ns, not significant.

| Phenotype/Genotype | BNS |  |  |  |  | LCNS |  |  |  |  |
| --- | --- | --- | --- | --- | --- | --- | --- | --- | --- | --- |
|  | WT | *glcat14a/b/d-1* | *p* | *glcat14a/b/d-2* | *p* | WT | *glcat14a/b/d-1* | *p* | *glcat14a/b/d-2* | *p* |
| Flowering time (days) | 24.1 ± 0.3 | 26.2 ± 0.4 | <0.0002 | 25.0 ± 0.5 | ns | 22.7 ± 0.3 | 24.8 ± 0.4 | <0.0021 | 25.1 ± 0.3 | <0.0002 |
| Stem length (mm) | 22.9 ± 0.6 | 16.0 ± 0.8 | <0.0001 | 15.5 ± 0.7 | <0.0001 | 22.3 ± 0.6 | 10.4 ± 0.4 | <0.0001 | 10.7 ± 0.4 | <0.0001 |
| Silique length (mm) | 12.2 ± 0.2 | 11.4 ± 0.4 | ns | 11.1 ± 0.4 | ns | 10.9 ± 0.7 | 4.3 ± 0.7 | <0.0001 | 4.7 ± 0.6 | <0.0001 |
| Seed set (%) | 80.5 ± 4.0 | 59.8 ± 2.6 | <0.0332 | 69.6 ± 6.0 | ns | 72.6 ± 9.8 | 6.3 ± 3.9 | <0.0001 | 6.1 ± 3.8 | <0.0001 |
| Viable PGs (%) | 99.8 ± 0.1 | 61.1 ± 14.3 | <0.0332 | 62.1 ± 9.6 | <0.0332 | 98.3 ± 0.9 | 23.9 ± 5.0 | <0.0001 | 21.8 ± 10.6 | <0.0001 |
| Aborted PGs (%) | 0.2 ± 0.1 | 38.9 ± 14.3 | <0.0332 | 37.9 ± 9.6 | <0.0332 | 1.7 ± 0.9 | 76.1 ± 5.0 | <0.0001 | 78.2 ± 10.6 | <0.0001 |
| Pollen germination (%) | 72.5 ± 3.4 | 36.0 ± 1.0 | <0.0332 | 40.3 ± 0.4 | <0.0332 | 63.3 ± 2.2 | 16.9 ± 6.8 | <0.0001 | 20.3 ± 8.2 | <0.0001 |
| PT length (μm) | 247.4 ± 27.6 | 173.4 ± 18.5 | <0.0332 | 166.1 ± 5.6 | <0.0332 | 241.1 ± 17.7 | 130.7 ± 2.1 | <0.0021 | 135.0 ± 9.3 | <0.0021 |

**Supplementary Table 3.**
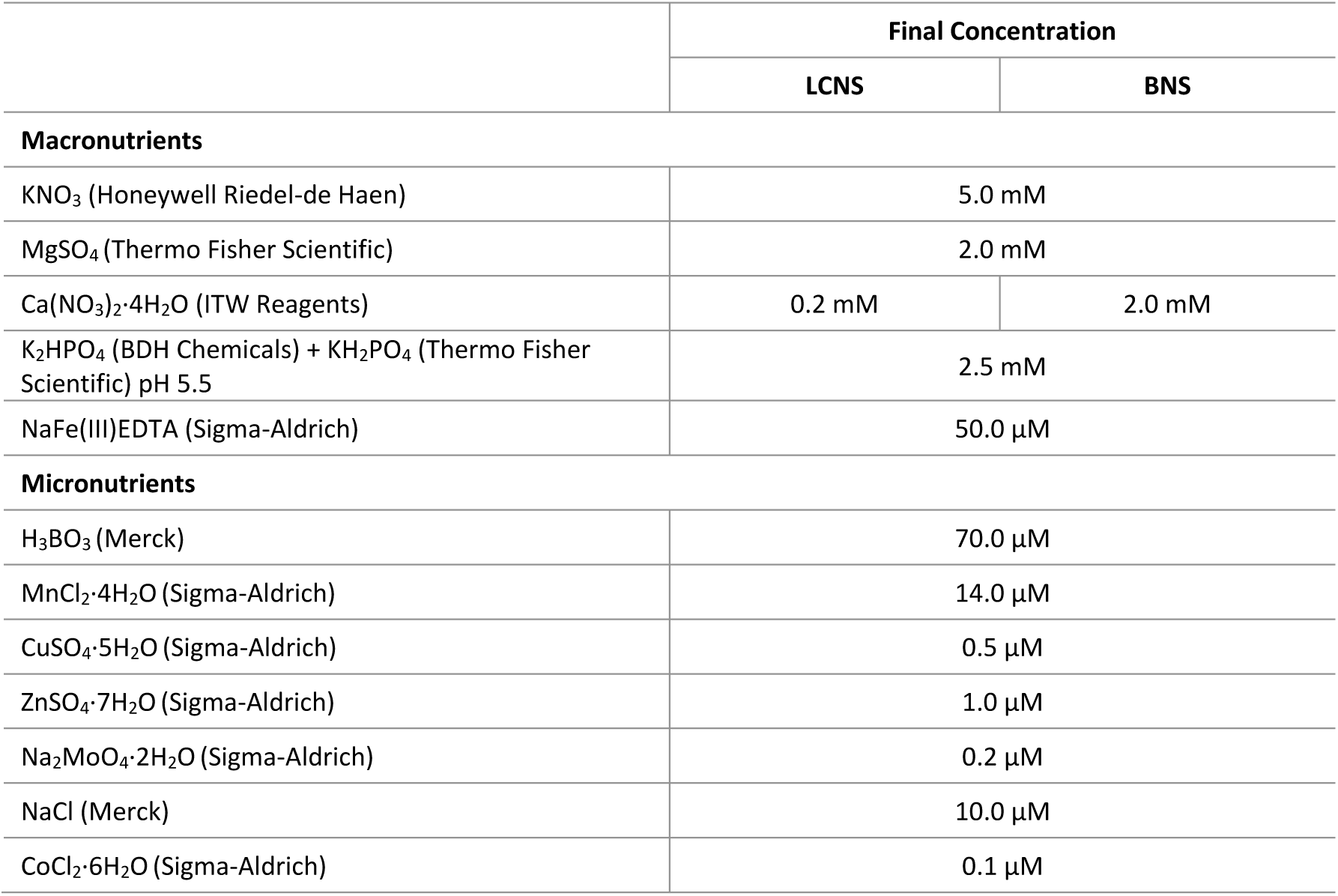
Nutrient solution composition.

**Supplementary Table 4.**
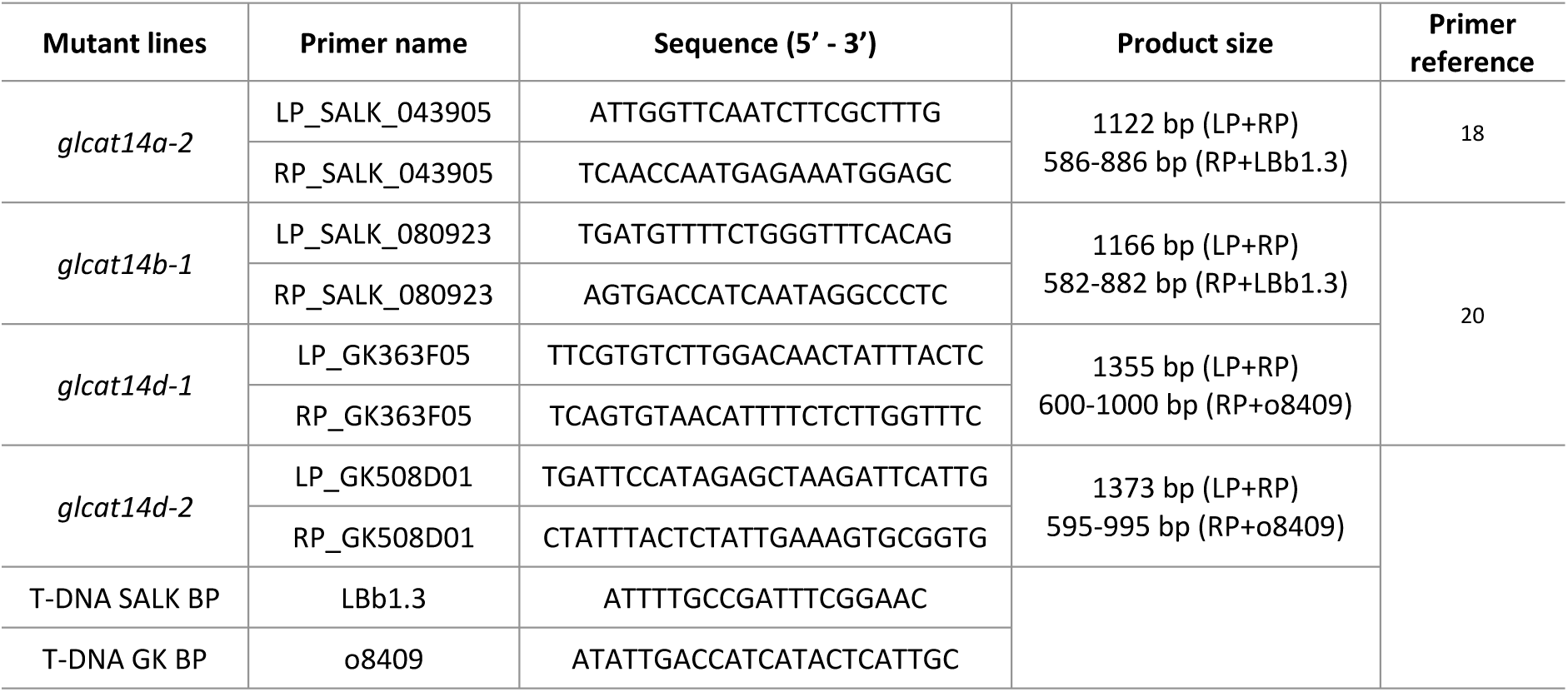
List of primers used to genotype T-DNA insertion lines. Abbreviations: bp, base pair; BP, T-DNA border primer; LP, left genomic primer; RP, right genomic primer.

**Supplementary Table 5.**
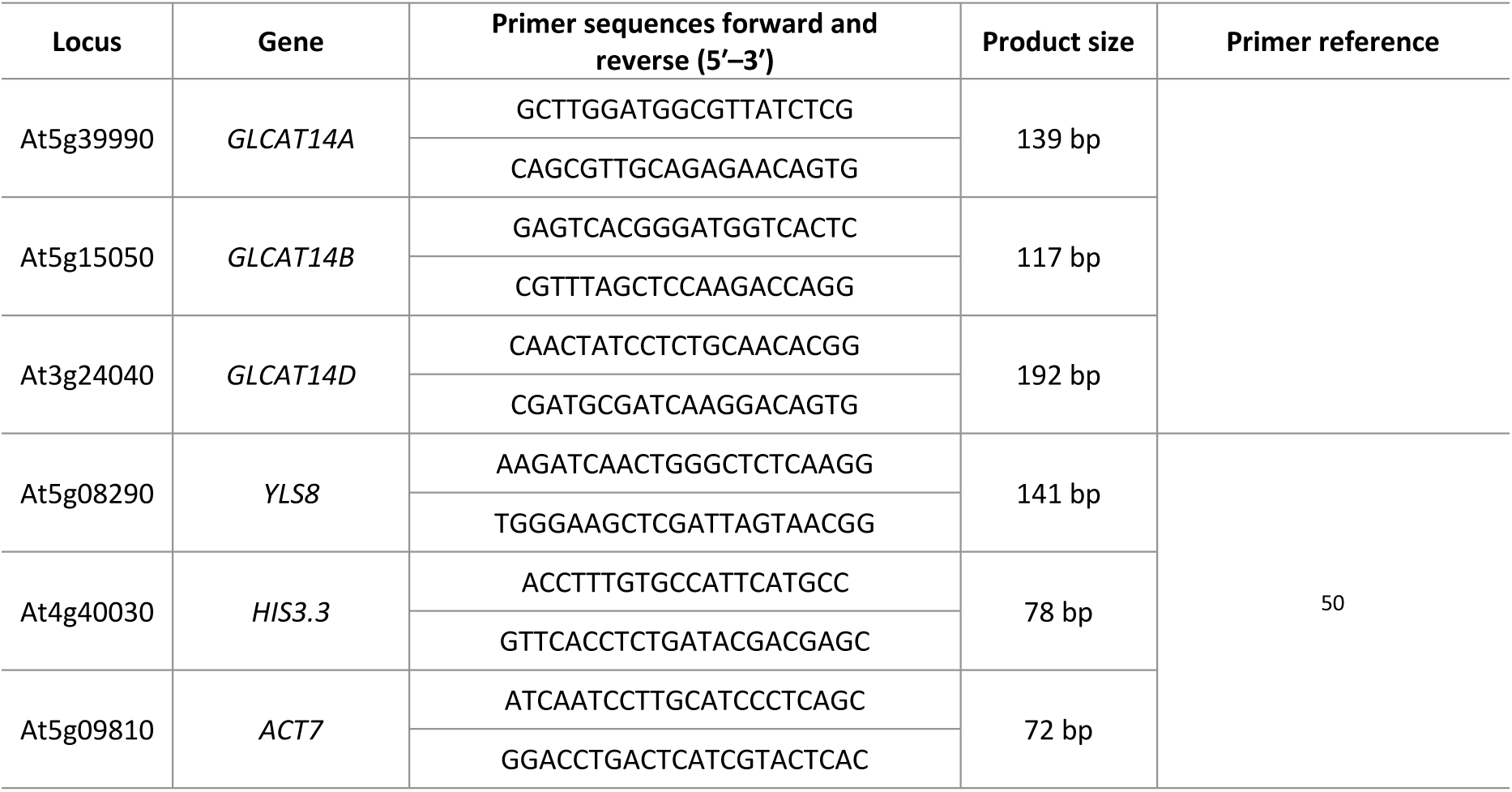
List of genes and primer sequences used for qPCR analysis. Abbreviations: bp, base pair.

**Supplementary Table 6.** Amplification efficiencies, correlation coefficients (R^2^), slope, and melting temperature of qPCR primers of target genes.

| Gene | Efficiency (%) | $R^2$ | Slope | Melting temperature (°C) |
| --- | --- | --- | --- | --- |
| *GLCAT14A* | 95.9 | 0.995 | -3,423 | 80.5 |
| *GLCAT14B* | 95.2 | 0.986 | -3.441 | 83.5 |
| *GLCAT14D* | 102.4 | 0.986 | -3.266 | 82 |

